# Fungal spore seasons advanced across the US over two decades of climate change

**DOI:** 10.1101/2024.10.21.619107

**Authors:** Ruoyu Wu, Yiluan Song, Jennifer R. Head, Daniel S. W. Katz, Kabir G. Peay, Kerby Shedden, Kai Zhu

**Affiliations:** School for Environment and Sustainability, University of Michigan, Ann Arbor, MI 48109, USA; Institute for Global Change Biology, University of Michigan, Ann Arbor, MI 48109, USA; Michigan Institute for Data and AI in Society, University of Michigan, Ann Arbor, MI 48109, USA; Department of Epidemiology, University of Michigan, Ann Arbor, MI 48109, USA; School of Integrative Plant Science, Cornell University, Ithaca, NY 14853, USA; Department of Biology, Stanford University, Stanford, CA 94305, USA; Department of Earth System Science, Stanford University, Stanford, CA 94305, USA; Department of Statistics, University of Michigan, Ann Arbor, MI 48109, USA

**Keywords:** Airborne allergen, Allergy, Climate change fingerprint, Human pathogen, Phenology, Public health

## Abstract

Phenological shifts due to climate change have been extensively studied in plants and animals. Yet, the responses of fungal spores—crucial organisms that play important roles in ecosystems and act as airborne allergens—remain understudied. This knowledge gap in global change biology hinders our understanding of its ecological and public health implications. To bridge this gap, we acquired a long-term (2003 ∼ 2022), large-scale (the continental US) dataset of airborne fungal spores collected by the US National Allergy Bureau. We first pre-processed the spore data by gap-filling and smoothing. Afterward, we extracted ten metrics describing the phenology (e.g., start and end of season) and intensity (e.g., peak concentration and integral) of fungal spore seasons. These metrics were derived using two complementary but not mutually exclusive approaches—ecological and public health approaches, defined as percentiles of total spore concentration and allergenic thresholds of spore concentration, respectively. Using linear mixed effects models, we quantified annual temporal shifts in these metrics across the continental US. We revealed a significant advancement in the onset of the spore seasons defined in both ecological (11 days, 95% confidence interval: 0.4 ∼ 23 days) and public health (22 days, 6 ∼ 38 days) approaches over two decades. Nevertheless, the total spore concentration in an annual cycle and in a spore allergy season tended to decrease over time. The earlier start of the spore season was significantly correlated with climatic variables, such as warmer temperatures and altered precipitations. Overall, our findings suggest possible climate-driven advanced fungal spore seasons, highlighting the importance of climate change mitigation and adaptation in public health decision-making.

## 1. Introduction

The shifts of phenology—the timing of seasonal activities of living organisms—in response to climate change are evident and extensively documented across diverse ecosystems, from polar lands to tropical oceans (Bradley et al., 1999; Menzel, 2000; Parmesan, 2007; Walther et al., 2002). Most phenological research focuses on conspicuous macroorganisms, which have gained significant public attention. For instance, North American and European studies show a consistent trend towards earlier biological spring events, such as bird migrations (Sparks, 1999), butterfly appearance (Roy & Sparks, 2000), amphibian spawning (Beebee, 1995; Kusano & Inoue, 2008), and plant leafing and flowering (Menzel et al., 2020; Piao et al., 2019), in response to a warming climate. However, corresponding shifts in microorganism seasonality, such as the phenology and intensity of fungal spore production, have received limited attention.

Fungal spores represent an important life stage of fungi, which are key components of ecosystems; they are also major airborne allergens affecting public health. From an ecological perspective, fungal spores are reproductive propagules that enable fungi to reproduce and spread (Wyatt et al., 2013). Spores are often dispersed by air, water, or other organisms, facilitating the colonization of new environments (Wyatt et al., 2013). Variation in the quantity, composition, and arrival time of fungal spores has been demonstrated to change multiple ecological processes (Dighton, 2018; Dighton & White, 2017), such as fungal community assembly (Leopold & Fukami, 2021; Peay, 2018; Peay et al., 2012), decomposition rate (Smith & Peay, 2021), nutrient cycling (Orwin et al., 2011; Read & Perez-Moreno, 2003), and plant productivity and competition (Levetin, 2016; Peay, 2018; Van Der Heijden et al., 2008). From a public health perspective, exposure to airborne fungal spores may trigger allergic reactions, resulting in asthma exacerbations or aggravating allergic rhinitis. The allergic symptoms to inhalation could range from mild respiratory discomfort to severe respiratory distress, sometimes inducing life-threatening reactions (Horner et al., 1995; Hughes et al., 2022; Jenkins et al., 1981; Kurup et al., 2000). In asthmatic patients, especially children, dominant allergenic fungi, such as *Alternaria*, *Cladosporium*, *Aspergillus*, and *Penicillium*, can precipitate acute symptoms (Batra et al., 2022; Chen et al., 2014; Tham et al., 2017). A serum-specific IgE test in the US suggests that 22% of allergic patients have allergic responses to at least one fungal allergen (Kwong et al., 2023). Furthermore, hospital admissions and mortalities related to asthma increase on days marked by high fungal spore concentrations (R. W. Atkinson et al., 2006; Dales et al., 2000; Newson et al., 2000). Due to their importance for both ecological communities and public health, understanding the phenological shifts of fungal spores is crucial for assessing the impact of climate change on ecosystems and informing decision-making in public health.

Fungal physiological studies suggest that climate change has the potential to influence the phenology and intensity of elevated airborne fungal spore concentrations in a year (herein referred to as fungal spore seasons) through various mechanisms. Both temperature and precipitation have been shown to affect fungal life cycles and the presence of spores in the atmosphere, though these effects are highly species-dependent and may vary in direction depending on the specific taxon (Diez et al., 2013; Gregory, 1967; Tilak, 2009). In the early stages of fungal growth, elevated temperatures can stimulate the rapid growth of some fungi species and induce their subsequent spore production (Jesús Aira et al., 2012); the post-rainfall moisture retained in the soil and leaves can provide the necessary substrates for fungal development and subsequent spore production in some species (Ganthaler & Mayr, 2015). A previous study on mushroom fruiting phenology suggested a widening fruiting season in Europe (Kauserud et al., 2012). Subsequently, the process of spore release into the air is also influenced by environmental conditions such as warmth, humidity, or dryness, which facilitate the detachment of spores from their parent structures (Gabey et al., 2010; Katial et al., 1997; Tilak, 2009), leading to an upsurge in airborne spore density (Ganthaler & Mayr, 2015). However, the process of aerosolization—the wide dispersion of spores through the air—tends to be more efficient under drier conditions. Dryness can reduce the agglutination of spores, thereby increasing their potential to become airborne and travel over longer distances (Sache, 2000). The ultimate viability of spores can adapt to various climatic conditions, with some spore taxa surviving across wide environmental ranges (Wyatt et al., 2013), and others strictly requiring certain temperatures and humidity levels (Golan et al., 2023). Apart from impacting fungi physiology directly, climate change could also alter spore phenology indirectly by interacting with other global change factors, such as land use land cover change (Delmas et al., 2024), and plant phenological change (Dickie et al., 2010). This multistage perspective underscores the nuanced impacts of climate patterns on fungal spore dynamics, thus impacting airborne fungal spore phenology and intensity. In return, this could have implications for ecosystems and public health. Therefore, it is critical to closely monitor and analyze the temporal patterns of fungal spores as they respond to ongoing climatic shifts.

Despite the recognized importance of fungal spores and potential shifts in their seasonality under a changing climate, comprehensive evaluations are limited. Several local studies in Europe reported inconsistent long-term trends of fungal spore phenology and intensity. For example, under similar rising temperatures, some studies reported an earlier spore season (Damialis et al., 2015; Grinn-Gofroń et al., 2016; Lam et al., 2024; Ščevková et al., 2016), while others suggested a delayed season (Damialis et al., 2015). Likewise, the trends in peak concentration were also heterogeneous over space in Europe (Damialis et al., 2015; Millington & Corden, 2005). Such inconsistencies could be attributed to differences in climate regime and fungal spore communities among local studies, usually constrained to one or two locations. An analysis synthesizing long-term trends from multiple locations could help provide generalizable insights on the overall change of fungal spore seasons on a large scale.

Fungal spore seasons can be characterized using diverse approaches, motivated by ecological dynamics and public health objectives. The start and end of spore seasons are commonly defined as the time when cumulative spore concentrations exceed a specific proportion of the annual total (Grinn-Gofroń et al., 2020; Katotomichelakis et al., 2016; Sadyś et al., 2016). This method identifies seasonal metrics based on the natural variations of fungal spores, providing understanding from an ecological perspective. Alternatively, the start and end of spore seasons are defined as the days when spore concentrations exceed and drop below critical thresholds that trigger health responses, such as allergic symptoms in susceptible populations (Clot, 2001; Giorato et al., 2000; Glick et al., 2021; Sánchez Mesa et al., 2003). This approach does not take into account the differences in total spore emission but focuses on the impacts of spore concentrations on human health, providing understanding from a public health perspective. While these approaches focus on different aspects—one on ecological dynamics and the other on public health—they are not mutually exclusive. In fact, the ecological cycles often align with the periods of heightened health risk, though the thresholds and sensitivities may vary. By integrating both the ecological cumulative integral method and the public health threshold method, we obtain a more comprehensive understanding of spore seasons that captures both their environmental and health-related significance.

In this study, we aim to quantify the long-term annual trends in the phenology and intensity of fungal spore seasons and their relationships with climate change. To achieve this, we analyze data from aerial samplings collected over 20 years (2003 ∼ 2022) from 55 monitoring stations across the continental US. Given the lack of studies on North American fungal spore seasonality, our study focuses on total spore concentration to fill this important gap. Our methodology integrates two approaches to define the spore seasons: the first uses cumulative integrals to define ecological spore seasons, consistent with prior ecological studies (herein: the ecological approach); and the second applies an allergy threshold estimated by a clinical study to identify spore allergy seasons (Caillaud et al., 2018), an approach consistent with current public health messaging and recommendations (herein: the public health approach). We subsequently employ statistical methods to quantify the trends in fungal spore season metrics and their relationships with climatic variables, including mean annual temperature and total annual precipitation. We hypothesize that the fungal spore phenology has shifted from 2003 to 2022, identified by an earlier start and longer length of the spore season. We also anticipate increases in the intensity of fungal spore seasons, such as peak and annual concentration, mirroring the shifts in allergenic pollen phenology in North America (Anderegg et al., 2021). Additionally, we hypothesize that the fungal spore season metrics correlate with climatic variables, including temperature and precipitation during the same period. Our study deepens the understanding of fungal spore ecology and could inform health-related policies and actions in response to the impact of climate change on allergenic fungal spores.

## 2. Material and methods

### 2.1. Data source for fungal spore concentrations and climate variables

We acquired a fungal spore concentration dataset from the National Allergy Bureau (NAB) counting stations, part of the American Academy of Allergy, Asthma & Immunology Aeroallergen Network (Table S1, Fig. S1). NAB has a long history of monitoring aeroallergens in the United States, thus providing long-term spore data over a large spatial scale (Levetin et al., 2022). To our knowledge, the NAB spore data is the most accurate spore data available because of their stringent sampling protocol (Portnoy et al., 2004). At each NAB station, airborne fungal spores are sampled over a 24-h period, primarily using a Burkard spore trap, though some stations still employ Rotorod samplers (Portnoy et al., 2004). According to the guidelines for setting up an NAB station, the sampler is positioned on a rooftop free of obstructions, with no unusual local sources of fungal spores surrounding it (Portnoy et al., 2004). Finally, the collected spores are identified and counted by certified experts, whose certifications need to be maintained (Portnoy et al., 2004). Daily average fungal spore concentrations (spores m^-3^) are calculated by the number of spores measured divided by the volume of air sampled over 24 hours. While sampling protocols, e.g., sampling devices, may differ across certain sites, our focus on temporal trends in fungal spore phenology is not likely to be affected by these spatial variations. In the subsequent statistical analyses, we also incorporated site-specific random effects in the linear mixed-effects models, helping to account for any potential variations introduced by differences in sampling across stations. Fungal spores were classified to the genus level whenever possible, with Cladosporiaceae and unidentified spores accounting for 44% and 46% of the total concentration, respectively (Fig. S2). To include the large proportion of unidentified spores and to gain a comprehensive understanding of overall spore seasonality, we focused on the total concentration of all fungal spore taxa in our analysis. Importantly, we did not detect significant temporal changes in the proportions of these two dominant taxa over the study period, supporting the validity of focusing on total spore concentration (Table S2).

From the NAB dataset, we retrieved a time series of total fungal spore concentration from 2003 to 2022 from 75 counting stations across the continental US. For robustness in extracting fungal spore season metrics, we selected stations that reported non-zero concentrations in at least ten days per year for at least three years. Overall, 638 station-year combinations across 55 NAB counting stations met our inclusion criteria. These stations are located in eight ecoregions in the continental US: 28 stations in the Eastern Temperate Forests ecoregion, 13 in the Great Plains ecoregion, six in the Mediterranean California ecoregion, and four in the North American Deserts ecoregion (Fig. S1; US EPA, 2015). The stations span a wide climatic gradient, with mean annual temperature ranging from −11.7 °C to 41.6 °C, and total annual precipitation ranging from 0 mm to 2,115.9 mm.

We obtained the climate data at each station from the National Aeronautics and Space Administration (NASA)-Daymet project (Thornton et al., 2022). The Daymet project provides 1 km gridded daily surface climate data, covering the whole duration and spatial coverage of the NAB spore data. Previous study has used the Daymet dataset for analyzing NAB data (Katz et al., 2023). We calculated mean annual temperature (MAT, °C) and total annual precipitation (TAP, mm) at each station for the years with spore sampling.

### 2.2. Pre-processing of spore data

Most of the station-year combinations contained missing data (Table S1; Figs. S3 and S4). Some smaller gaps occurred due to stations not operating during weekends. Larger gaps existed in winter seasons at a few stations, as these stations are mainly set up for pollen monitoring and might suspend monitoring during winter when pollen is not present. The missing data caused information loss and introduced potential uncertainty and bias in calculating spore season metrics, such as the start and end dates of the spore allergy season. For instance, the actual start date of the spore allergy season could not be accurately detected if it falls within a gap in data collection. Note that true zeros (no spore detected) are considered valid data points while missing data refers to instances where no measurement was recorded due to gaps in data collection or station inactivity. We mitigate issues associated with missing data and improve the accuracy of subsequent analyses in two ways. First, small gaps in each station, which were fewer than 14 consecutive days, were linearly interpolated by taking the first data before and after the gap, a method commonly used to handle missing data in aerobiological studies (Table S1; Anderegg et al., 2021; Gabarra et al., 2002; Picornell et al., 2021). Second, station-year combinations with large gaps were excluded from our analysis, and the detailed criterion will be stated for each fungal spore season metric subsequently.

To mitigate the impact of observational errors and day-to-day variability in fungal spore concentration, we employed the Whittaker-Henderson smoothing method for each station (Whittaker, 1922). The Whittaker-Henderson smoothing has been widely adopted in prior phenological research (P. M. Atkinson et al., 2012; Li et al., 2020). We used a smoothness parameter of 100 for the Whittaker-Henderson smoothing. To ensure the robustness of our results, we conducted a sensitivity analysis by varying the smoothness parameter, lambda, from ten to 500 (Fig. S5). We then tested whether the estimated temporal trends in fungal spore season metrics remained consistent across different values of lambda using linear regression (*t*-test).

This sensitivity analysis allowed us to evaluate the influence of the smoothing parameter on our results, ensuring that our approach was robust across a broad range of smoothing parameters. The pre-processed data were subsequently used to extract fungal spore metrics.

### 2.3. Characterizing fungal spore season

#### 2.3.1. Detecting seasonal cycles

To characterize the fungal spore season in each station, we visualized fungal spore calendars by averaging raw daily concentrations (without gap-filling and smoothing) from 2003 to 2022 (Fig. S4). Because the seasonal cycles of spore concentration did not coincide with calendar years (i.e., January 1st ∼ December 31st) in many stations, we defined spore years to calculate meaningful summary metrics (e.g., start and end of season) (Fig. 1a; Fig. S4). Specifically, we computed the median fungal spore concentration of each day for years with available data in each station; the beginning day of a spore year was then defined as the day when the trough value appeared in each station’s spore median concentration curve (Fig. 1a). After pre-processing raw data and defining the spore year for each station, we assessed the data completeness for each station and spore year combination by dividing the number of days with available data by 365. The data completeness subsequently served as a threshold criterion for the inclusion of station-spore year combinations in our future analysis.

**Fig. 1.**
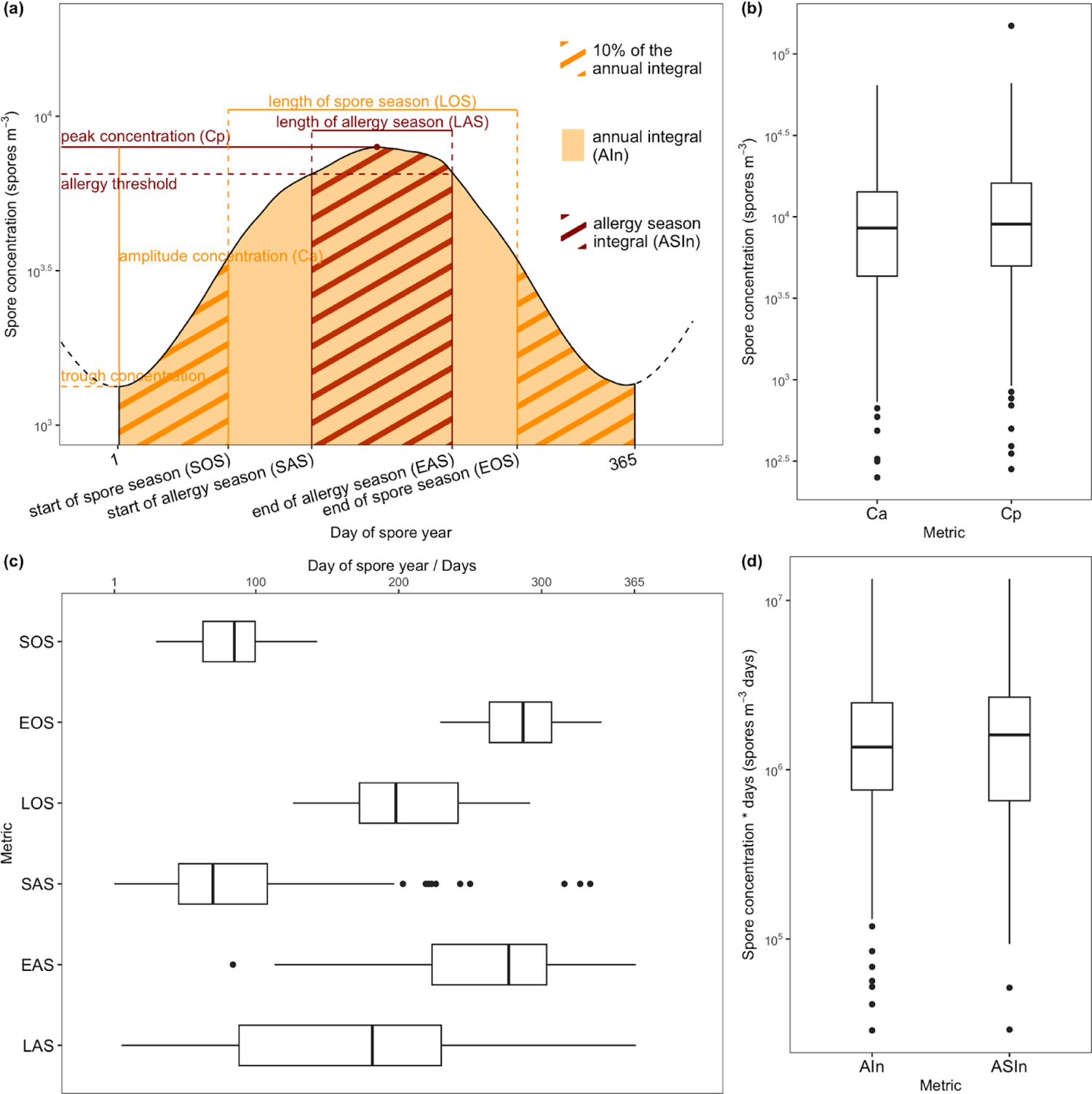
Definition and numerical summary of ten fungal spore metrics. (a) The annotation of ten fungal spore metrics. Orange indicates metrics defined in the ecological approach, while dark red indicates metrics defined in the public health approach. Each box represents each metric’s interquartile range (IQR), with the median value indicated by the line inside. Whiskers extend to the minimum and maximum values within 1.5 times the IQR, while points beyond the whiskers are considered outliers. (c) Phenology metrics are in the unit of day of spore year (DOY) or days (for season lengths LOS, LAS), while intensity metrics are in the unit of (b) spores m^-3^ or (d) spores m^-3^ days, displayed on a natural logarithm-transformed scale. The abbreviations of metrics used in (b), (c), and (d) are defined in (a).

#### 2.3.2. Defining spore season phenology

Within each spore year, we extracted ten fungal spore season metrics, encompassing the phenology and intensity of fungal spore season, using both ecological and public health approaches (Fig. 1a). From an ecological approach, we characterized the start of spore season (SOS) and the end of spore season (EOS) as the day of the spore year (DOY) when the cumulative sum of smoothed daily spore concentrations exceeded the 10th and 90th percentile of the annual spore concentration integral, respectively (Fig. 1a). This definition allowed us to characterize the timings of seasonal increase and decrease of fungal spore concentration. We then calculated the length of spore season (LOS, days) by subtracting SOS from EOS (Fig. 1a). Since the annual spore integral was a principal reference in defining the spore season, we required the data completeness after interpolation of each station-spore year to be 100% for metric SOS, EOS, and LOS to ensure the accuracy of the values.

From a public health approach, we adopted 4,506 spores m^-3^ as the allergy threshold in our definition of allergy season. This value was computed by converting the relative risk threshold of Cladosporium mold concentrations, which were linked to increases in prescribed allergy medication sales (Caillaud et al., 2018), to a total spore concentration based on the relative abundance of Cladosporium in our dataset (Fig. S2). Acknowledging that relying on the threshold from a single study could limit the generalizability of our results, we tested the robustness of the allergy threshold using a different NAB-defined moderate allergy risk level of 6,500 spores m^-3^ (Portnoy et al., 2004). Detailed methods to acquire the threshold and robustness test can be found in the supplementary text. Therefore, the start of allergy season (SAS) and the end of allergy season (EAS) is the first and last day of the spore year (DOY) of the first and last ten consecutive days with the smoothed spore concentration above the threshold of 4,506 spores m^-3^ (Fig. 1a). Since sampling may not cover a whole spore year, it is possible for sampling to start after the true SAS or end before the true EAS. To address this issue, we excluded detected SAS if ten consecutive days before this SAS contained missing data. Likewise, the detected EAS was removed if any of the ten consecutive days following it were incomplete. This method ensures that both SAS and EAS are accurately represented and not influenced by gaps in data, ultimately improving the reliability of the extracted metrics. The length of allergy season (LAS, days) was the interval between SAS and EAS when both SAS and EAS were available (Fig. 1a).

#### 2.3.3. Defining spore season intensity

We also extracted metrics related to the intensity of fungal spore season using both ecological and public health approaches. For each station-spore year, we calculated the peak and trough concentrations by taking the maximum and minimum spore concentrations in a station-spore year, respectively (Fig. 1a). Therefore, the amplitude concentration (Ca, spores m⁻^3^) was defined by the difference between peak concentration and trough concentration (Fig. 1a). We calculated the total spore concentration in an ecological spore year as the annual integral (AIn, spores m^-3^ days) by multiplying the average daily concentration by 365 days (Fig. 1a). From a public health approach, the peak concentration (Cp, spores m⁻^3^) was defined as the maximum spore concentration in a station-spore year (Fig. 1a). We also defined total spore concentration in a spore allergy season as the allergy season integral (ASIn, spores m^-3^ days) by multiplying the average daily concentration during the allergy season by the LAS (Fig. 1a). We adopted the following filtering criteria on data completeness for each station-spore year to balance the quality of metric extraction and sample size (Tables S3 and S4). We kept station-spore year combinations with more than 80% available data after interpolation for metrics Ca, Cp, and AIn; We kept station-spore year combinations with more than 80% available data after interpolation during the allergy season for metric ASIn. After extracting all the metrics, we characterized the overall spore season phenology and intensity across all the stations by taking the median value of each metric across all years in the study period.

### 2.4. Detecting temporal trends

We quantified the temporal trends of the ten fungal spore metrics at the continental US level. To focus on the long-term trends of the ten metrics, we included stations with five or more years of extracted metrics. To reduce data skewness, we applied a natural log transformation to all the intensity metrics in our analysis, specifically Ca, Cp, AIn, and ASIn. To explore the trends in metrics at each station, we used Theil-Sen linear regression of each metric against year at each station, a nonparametric method robust to outliers and suitable for exploratory data analysis with noisy metric values (Fig. S6; Sen, 1968; Theil, 1950).

We employed linear mixed-effects models to capture the overall temporal trends in spore metrics across all stations. Instead of assuming independence among all data points, the linear mixed-effects model accounts for within-station correlation and between-station variability, allowing for a more accurate analysis of our nested data. We fitted the following linear mixed-effects model of each fungal spore metric against year, with random intercepts and slopes on stations:

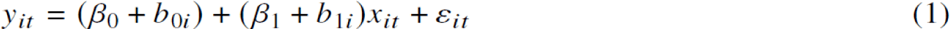

where the response *y_it_* is each fungal spore metric (with intensity metrics Ca, Cp, AIn, and ASIn log-transformed) in each station *i*, year *t*, while the independent variable *x_it_* here is year *t*. A fixed intercept *β*_0_ indicates the average spore metric across all stations when *x_it_* is zero, *b*_0*i*_ is the random intercept for station *i*, capturing the station-specific deviation from the overall average when *x_it_*is zero, the fixed slope, *β*_1_, quantifies the average temporal trend in the fungal spore season metric per year across all stations, *b*_1*i*_ is the random slope for station *i*, accounting for the variability in the rate of change specific to that station, random error *ε_it_* is assumed to be independent and normally distributed with mean 0 and variance *σ^2^*.

We then used the model-estimated parameters and predicted random effects to predict both overall trends and station-specific trends:

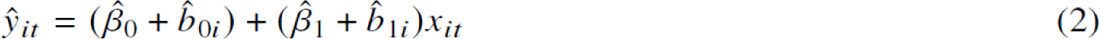

where *ŷ* is the predicted value of the fungal spore season metric at station *i* in year *t*. β^^^_0_ and β^^^_1_ are the point estimates of the fixed intercept and slope, representing the overall average across all stations. Similarly, *b*^^^_0i_ and *b*^^^_1i_ are the point estimates of the random intercept and slope for station *i*, capturing the station-specific deviations from the overall trend. Therefore, the term (β^^^_1_ + β^^^_0_ + *b*^^^_0i_) predicts the fungal spore season metric at station *i* when *x* is zero, and the term (β^^^_1_ + *b*^^^_1i_) predicts the temporal trend in the spore metric at station *i* per year, *x* is the year.

### 2.5. Assessing climate change impact

The varying rates of climate change across the continental US, including divergent patterns of warming and drying (Portmann et al., 2009), may alter fungal spore season phenology and intensity in different ways across different areas. To investigate this, we analyzed the relationship between climate and observed metrics at each station. For each station-spore year, we averaged daily mean temperature (°C) to calculate the mean annual temperature (MAT, °C). Total annual precipitation (TAP, mm) was derived by integrating daily precipitation (mm) within each spore year. To analyze the effect of climate on spore season metrics, we fitted linear mixed-effects models sharing the same structure with the model (1), except using MAT*_it_* (°C) or TAP*_it_* (mm) at station *i* in year *t* as the independent variable *x_it_*, respectively. The fixed slope, *β*_1_, represents the average sensitivity of the spore metric to changes in climatic variables across all stations, and *b*_1*i*_ is the random slope for station *i*, accounting for the variability in the sensitivity of the fungal spore season metric specific to that station. Using equation (2) above, we predicted the fungal spore season metric under the climatic condition *x_it_* for each station and time point (Fig. S7). All computations were carried out using R 4.3.3 (Bates et al., 2015; Pinheiro & Bates, 1995; R Core Team, 2024).

## 3. Results

### 3.1. Fungal spore season characteristics

Spatial variations in spore seasonality, encompassing both phenology and intensity, were evident among different stations (Fig. 2; Fig. S4). Stations characterized by warmer, humid summers and cold, dry winters, such as Dayton, OH, Berkeley, MO, and Oklahoma City, OK, experienced an increase in airborne spore concentrations typically at the onset of the calendar year (Fig. 2a).

**Fig. 2.**
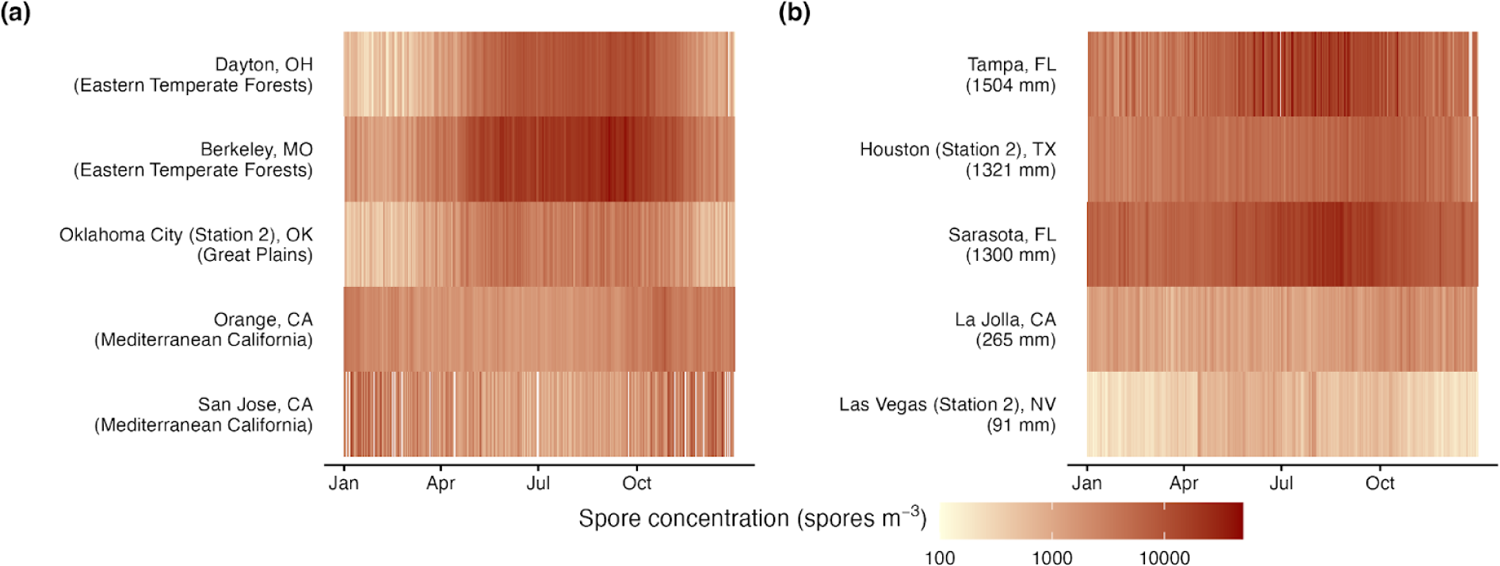
Fungal spore calendar for ten sampling stations. The daily long-term mean of fungal spore concentration is displayed, 2003-2022. Darker colors indicate higher concentrations while missing data are represented in white. (a) Stations show variable seasonality across ecological regions. (b) Stations with higher long-term mean of total annual precipitation (mm) show higher intensity of fungal spore seasons.

Conversely, stations situated within the Mediterranean California ecological region, known for its hot, dry summers and mild winters, displayed a different pattern, where winter and spring spore concentrations surpassed those observed in the summer. At these stations, a notable increase in spore concentrations began around the middle of the calendar year (Fig. 2a). The observed mismatch in seasonal patterns underscores the necessity for station-specific definitions of the “spore year” to account for the unique cycles at each location. The start of the spore year for each station can be found in the supplementary information (Fig. S4). Stations with distinct precipitations showed variability in spore season intensity. Stations with higher annual precipitation, such as Tampa, FL, Houston, TX, and Sarasota, FL, experienced higher spore concentrations throughout the year (Fig. 2b). Alternatively, stations receiving less precipitation, such as La Jolla, CA, and Las Vegas, NV, tended to record lower spore concentrations (Fig. 2b).

After defining the spore year for each station, we observed the overall spore season characteristics across the continental US. The ecological approach quantified the spore season as lasting a median of 198 days, with a median SOS of 85 day of spore year (DOY) and a median EOS of 287 DOY (Fig. 1c). From a public health approach, the allergy season started at 70 DOY and ended at 277 DOY, resulting in a median LAS of 182 days (Fig. 1c). The spore season intensity had a median amplitude concentration of 8,538 spores m^-3^ and a median peak of 9,009 spores m^-3^ (Fig. 1b). The median AIn and ASIn were 1.36 million spores m^-3^ days and 1.6 million spores m^-3^ days, respectively (Fig. 1d). The ecological and public health approaches lead to similar estimates of the phenology and intensity of spore seasons.

### 3.2. Temporal trends in spore metrics

Sixteen to 31 stations for each metric were included in our analysis (Table S3). We observed spatial variations in station-level trends of metrics. For instance, 20 out of 31 stations, including Twin Falls, ID, Oklahoma City, OK, and Tampa, FL, experienced an advancement in the start of the allergy season (SAS) over time (Fig. 3), while the SAS in other stations, such as Olean, NY, was delayed over time (Fig. 3). The annual spore integral in 15 out of 22 stations decreased over time, while seven stations were exposed to higher annual spore integral over time (Fig. S6). We also identified similar spatial heterogeneity in the trends of other metrics across different stations (Fig. S6).

**Fig. 3.**
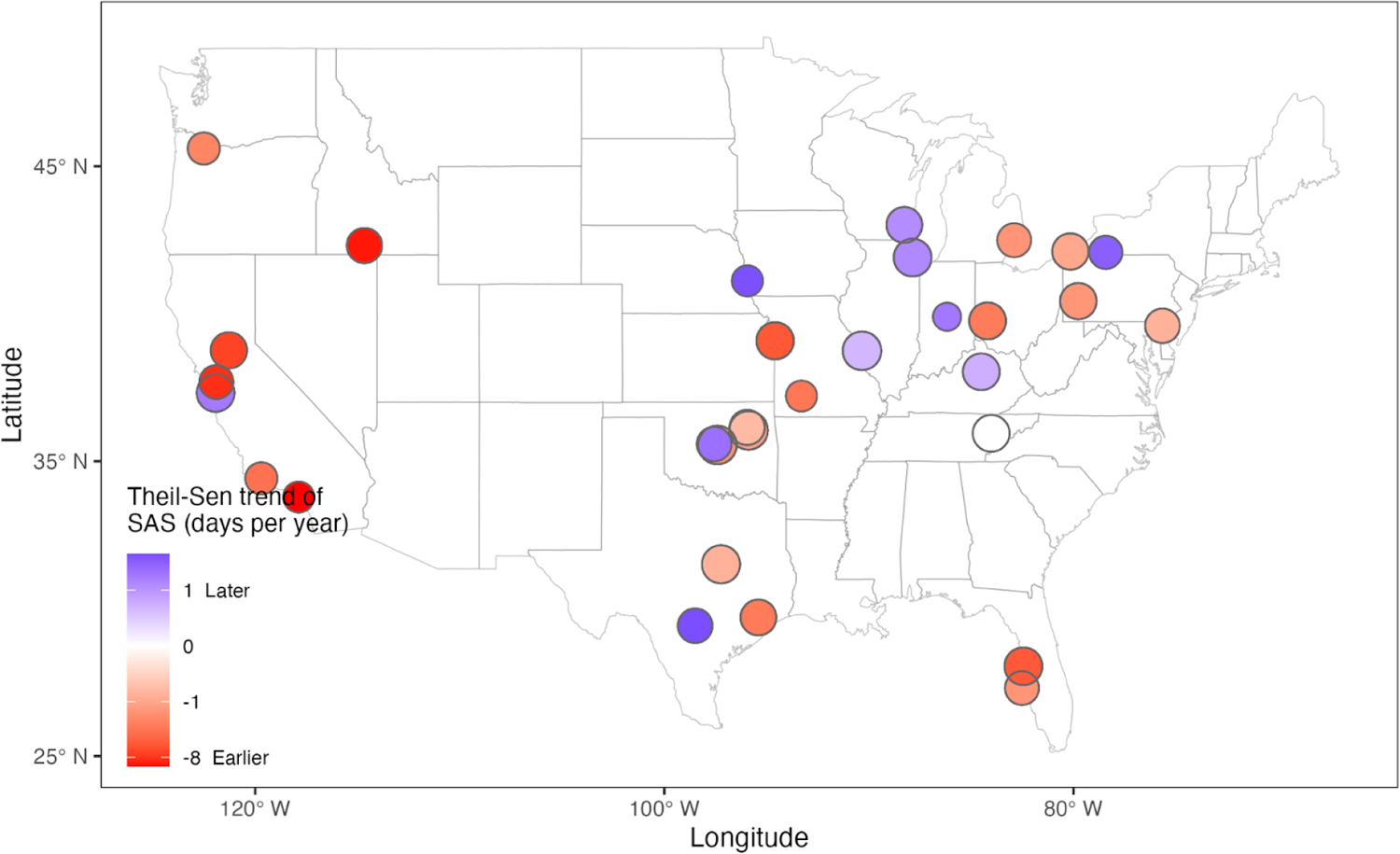
Station-level temporal trends of the start of allergy season (SAS). The trends were estimated by Theil-Sen linear regression. Warm colors indicate earlier seasons, and circle sizes are proportional to the years of data at each station.

To synthesize heterogeneous station-level temporal trends, we used linear mixed-effects models to evaluate the continental US-level trends. We quantified an overall significant advancing trend in the onset of spore seasons, using both the ecological and the public health approaches for calculating season start (SOS and SAS, respectively; Fig. 4a, f). Between 2003 and 2022, we detected an advancement in the SOS of 11 days, on average, across the continental US, corresponding to a rate of 0.59 (95% confidence interval: 0.019 ∼ 1.2) days year^-1^ (*p* < 0.05; Fig. 4a and Fig. 5; Table S3). The SAS in the continental US advanced significantly at the rate of 1.2 (0.3 ∼ 2) days year^-1^, equivalent to an advancement of 22 days from 2003 to 2022 (*p* < 0.05; Fig. 4f and Fig. 5; Table S3). In contrast, no significant shifts were found at the end of the spore season (EOS), while the end of the allergy season (EAS) became earlier significantly at the rate of 1.4 (0.37 ∼ 2.5) days year^-1^ over the two-decadal period (*p* = 0.998 for EOS, *p* < 0.05 for EAS; Fig. 4b, g and Fig. 5; Table S3). Length of the season changes were not significant, with LOS showing non-significant extensions of 11.6 days, occurring at a rate of 0.61 (−0.13 ∼ 1.4) days per year (*p* = 0.1; Fig. 4c and Fig. 5; Table S3); LAS getting shorter at a rate of 0.55 (−0.99 ∼ 2.1) days per year, resulting in ten days shorter insignificantly (*p* = 0.48; Fig. 4h and Fig. 5; Table S3). Regarding the intensity of the spore seasons, Ca and ASIn exhibited significant decline across the continental US over the study period (Fig. 4d, j), while Cp and AIn had non-significant declines (Fig. 4i, e). The detected continental US-level trends are robust to different smoothing parameter values and allergy thresholds.

**Fig. 4.**
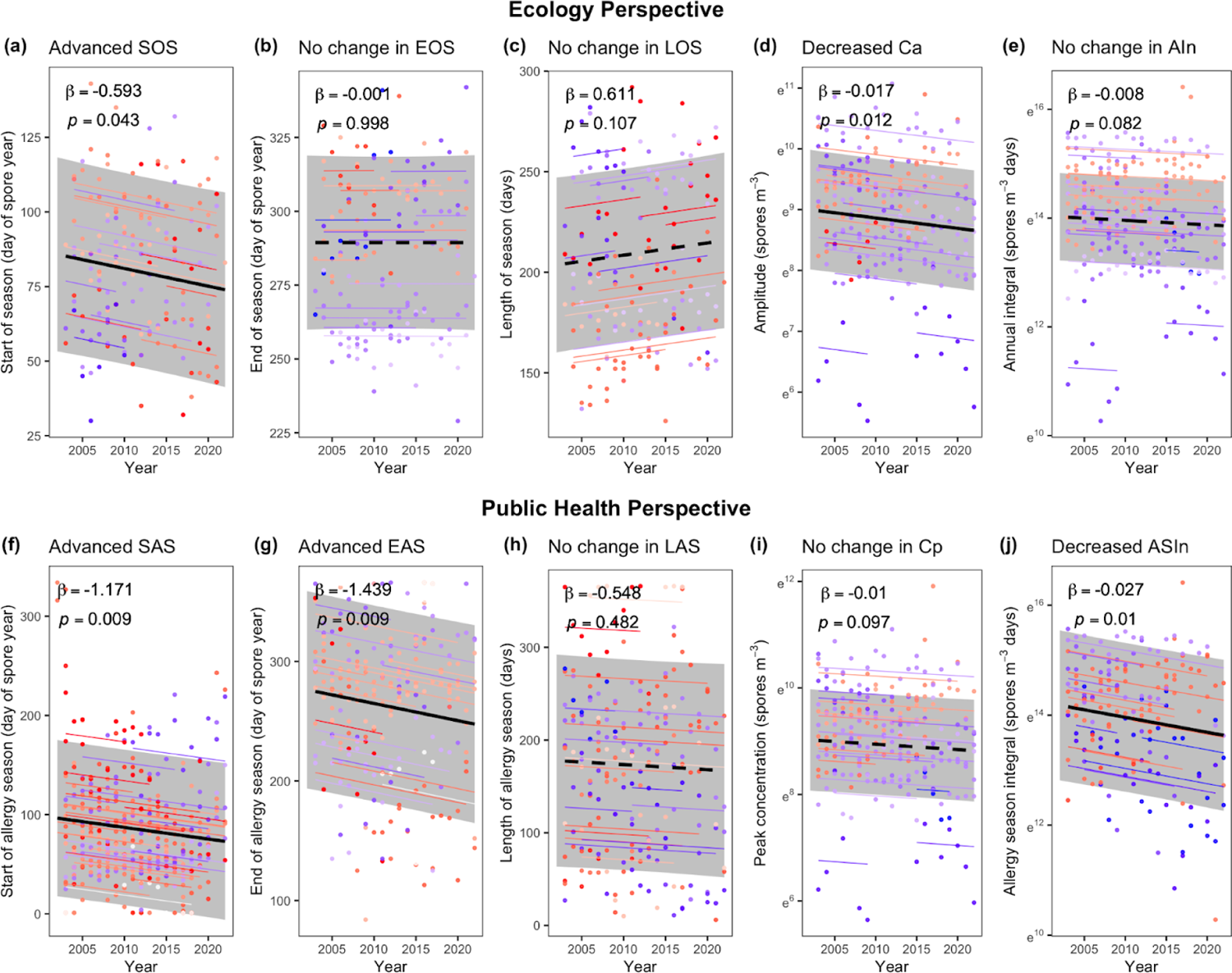
Temporal trends of ten fungal spore metrics across all the stations. The estimated trends from linear mixed effects models across stations of the year against (a) the start of season (SOS), (f) start of allergy season (SAS), (b) end of season (EOS), (g) end of allergy season (EAS), (c) length of season (LOS), (h) length of allergy season (LAS), (d) amplitude concentration (Ca), (i) peak concentration (Cp), (e) annual integral (AIn), and (j) allergy season integral (ASIn). Solid black lines indicate significant (*p* < 0.05, *t*-test) slopes. Shaded areas indicate the 95% confidence intervals of the fixed effect. Points are individual years at individual stations. Colors are station-level Theil-Sen linear regression slopes, with warmer colors indicating earlier days of year, longer days, or higher intensity. The slope of colored lines are model-predicted station-level trends. The top row presents metrics defined in the ecological approach, while the bottom row shows metrics defined in the public health approach. Intensity metrics are transformed using the natural logarithm.

**Fig. 5.**
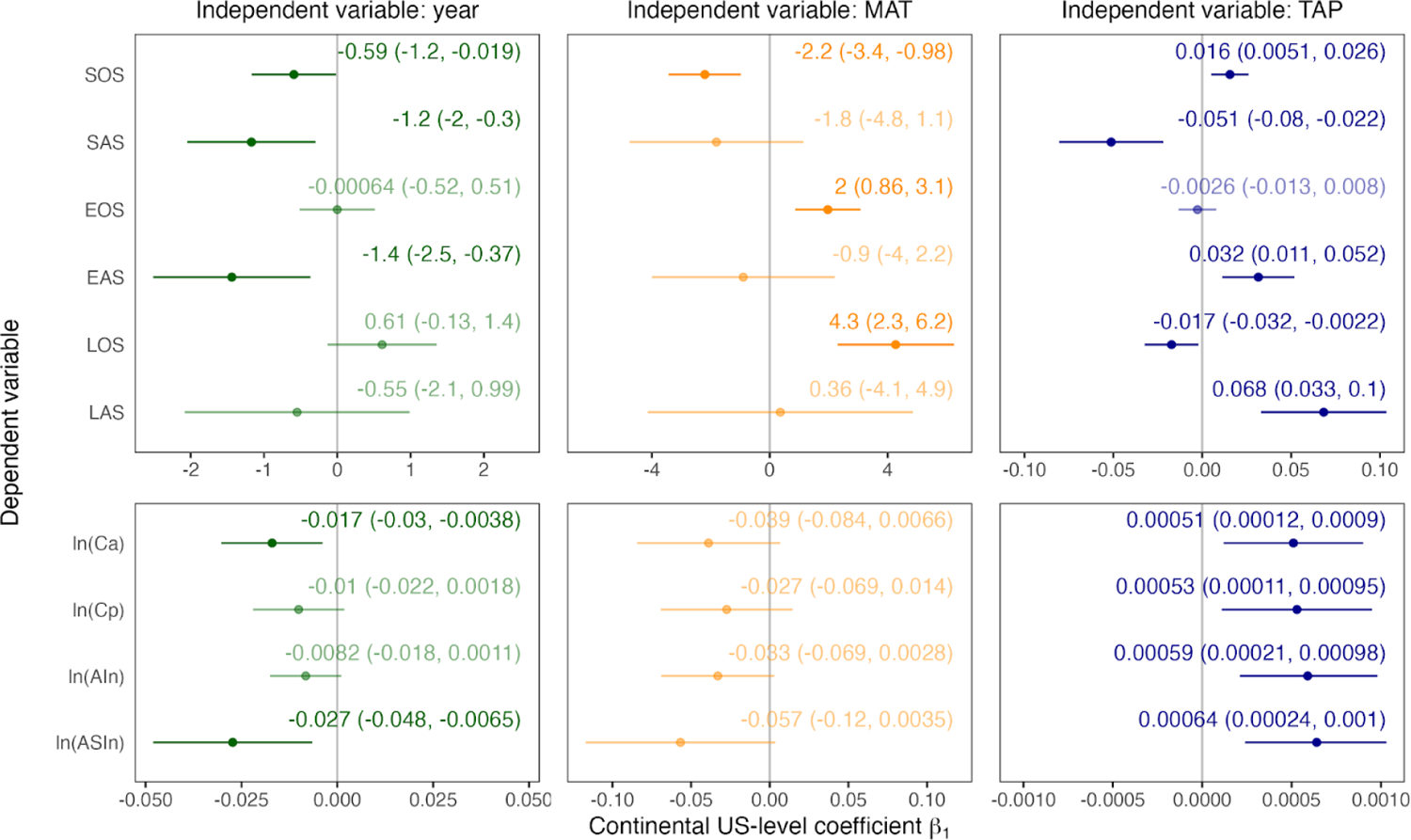
Summary of coefficients *β*_1_ in three mixed-effects models for each metric. Points are estimated values of *β*_1_. Error bars indicate the 95% confidence intervals of the fixed slope. Specific values are annotated. Colors are different independent variables in the model. Non-transparent colors present significant coefficients (*p* < 0.05, *t*-test).

Sensitivity analyses were conducted to ensure the robustness of our findings across different parameter settings and allergy thresholds. The sensitivity analysis on the smoothness parameter, lambda, for the Whittaker-Henderson smoothing, showed that both the SOS and SAS advanced significantly, regardless of the lambda value, which varied from ten to 500 (Fig. S5). Intensity metrics, including Ca, Cp, AIn, and ASIn, decreased consistently across the range of lambda values (Fig. S5). Additionally, using the NAB threshold of 6,500 spores m^-3^ for moderate allergy risk (Portnoy et al., 2004) in our analysis confirmed that both the ecological spore season and allergy season experienced significantly earlier onsets (Fig. S8). These sensitivity analyses reassure the consistency and robustness of our findings across different smoothing parameters and thresholds, supporting the conclusion that fungal spore seasons have experienced earlier starts over the last two decades, with a potential decline in intensity.

### 3.3. The relationship between spore season shifts and climate change

To further understand the long-term temporal trends, we analyzed the correlations between fungal spore season metrics and climate variables (Fig. 5, Fig. S8). On the one hand, MAT was significantly correlated with spore seasonality. Our findings indicate a significant negative correlation between temperature and SOS, implying that higher temperatures may be associated with an earlier onset of the spore season (*p* < 0.05; Fig. 5). Specifically, a 1 °C increase in MAT corresponded to advancement of the SOS by 2.2 days (95% confidence interval: 0.98 ∼ 3.4; Fig. 5). Later EOS was significantly related to warmer conditions: a 1 °C increase in MAT was correlated with a delay of 2 (0.86 ∼ 3.1) days in EOS (*p* < 0.05; Fig. 5). These result in a significantly positive correlation between LOS and temperatures, with an extension of 4.3 (2.3 ∼ 6.2) days per °C increase in MAT (*p* < 0.05; Fig. 5). On the other hand, TAP could significantly influence both spore seasonality and intensity. Rainfall had opposite impacts on the phenology of spore season and allergy season. Wetter years were correlated with later SOS and shorter LOS, with a delay of 0.016 (0.0051 ∼ 0.026) days in SOS and a shortening of 0.017 (0.0022 ∼ 0.032) days in LOS per mm increase in TAP (*p* < 0.05; Fig. 5). Conversely, more precipitation was characterized by earlier SAS and longer LAS (*p* < 0.05; Fig. 5). Numerically, a 1 mm increase in TAP was associated with 0.051 (0.022 ∼ 0.08) days of advancement in SAS and 0.068 (0.033 ∼ 0.1) days of extension of LAS (Fig. 5). Annual amplitude concentration, annual peak concentration, annual spore integral, and the allergy season integral were positively correlated with total annual precipitation (*p* < 0.05; Fig. 5). Overall, precipitation might have played a key role in altering fungal spore seasons due to its significant correlations with most season metrics from both ecological and public health approaches, along with an additional influence of temperature on some metrics.

## 4. Discussion

Our analysis supports the hypothesis of a multi-decadal, continental-scale shift in fungal spore phenology and intensity under climate change. Specifically, over the period from 2003 to 2022, across the continental US, we identified a significantly advancing start of fungal spore seasons—defined both by when fungal spore concentrations exceeded certain percentiles of total annual concentrations and by when fungal spore concentrations exceeded the clinical study-estimated allergy threshold. Additionally, our analysis shows a significant decrease in the amplitude concentration and total spore concentration in the allergy season, and non-significant decreases in the peak concentration and annual total spore concentration. Notably, these observed trends were found to be strongly correlated with climate change, particularly warming temperatures and altered precipitation. These results contribute to a growing body of evidence suggesting the impacts of climate change on phenological shifts of microorganisms, with implications for both ecology and public health.

### 4.1. Shifts in fungal spore seasonality

Our findings revealed notable shifts in fungal spore phenology across the continental US. Using two different, yet complementary, methods for determining seasons of elevated airborne spore concentrations, we documented a significant earlier onset of both the ecological spore season and the spore allergy season (Fig. 4a, f). Our observations aligned with published research from other regions. Notably, a study conducted over 19 years in Northern Italy highlighted a shift towards an earlier onset of the spore season by 0.7 days annually—findings that closely mirror our own (Marchesi, 2020). Studies from Poland and Slovakia reported more pronounced shifts, finding that the start of the spore season advanced four (Grinn-Gofroń et al., 2016) and 2.5 (Ščevková et al., 2016) days per year, respectively. On the other hand, contrasting patterns have been observed in other studies. A study in Greece noted a delayed spore season onset (Damialis et al., 2015), while research in Denmark recorded no clear trends in the phenology of the spore season (Olsen et al., 2020).

We observed significant decreases in total spore concentration in the allergy season, and non-significant decreases in peak concentration and annual total spore concentration over time (Fig. 4). The decreasing trends in peak spore concentrations and total annual concentration are consistent with findings from multiple studies (Damialis et al., 2015; Grinn-Gofroń et al., 2016; Marchesi, 2020; Olsen et al., 2020). Opposite trends in spore season intensity, such as increases in spore counts over time, have also been documented in long-term studies spanning 13 to 34 years in locations like Slovakia (Ščevková et al., 2016), the UK (Millington & Corden, 2005), and France (Sindt et al., 2016).

The inconsistencies across studies may arise from methodological differences. Many previous studies have centered their analysis on data from a single monitoring station, while our study draws from a larger geographic scope, incorporating data from multiple stations across the US. We also observed spatial variations in station-level trends of metrics (Fig. 3; Fig. S6). This could be due to regional differences in climate change trends between sites, local fungal spore composition, and other environmental factors, as well as high uncertainty in spore season metrics, making local trends in metrics different across sites. Our analysis of the continental US-level trends alleviates these problems to some extent by pooling data across sites, e.g., 16 stations for the start of spore season and 31 stations for the start of allergy season, increasing the generalizability of our conclusions. Our research emphasizes the need for comprehensive and methodologically consistent data collection across extensive geographic areas to fully understand spatial variations in fungal spore seasonality and its responses to environmental change.

The analysis of airborne fungal spores in our study contributes to the broader field of aerobiology, which also encompasses significant airborne allergens such as pollen. We can further contextualize our findings by comparing them with a recent study examining pollen trends across the continental US, which, like this study, also utilized the data from the National Allergy Bureau (NAB). This research documented a shift towards earlier pollen season onset, with start dates advancing by ∼20 days over the 28 years spanning 1990 to 2018 (Anderegg et al., 2021). These changes mirror the phenological shifts in fungal spore seasons observed in our research qualitatively and quantitatively, highlighting parallel trends in pollen and spore phenologies. However, in contrast to our results, an increase in the pollen annual integral was reported (Anderegg et al., 2021). Further contextualization in global change biology comes from comparing our research with phenological studies on macroorganisms like plants and insects.

Prior research has identified an overall trend towards earlier spring events across various taxonomic groups, with an average rate of 0.28 days per year (Parmesan, 2007). More specifically, tree phenology has been advancing by 0.33 days per year, while butterfly spring phenology has shown an advancement of 0.37 days per year (Parmesan, 2007). The rate at which these macroorganisms are experiencing spring advancements is in line with the advancement rate we have found in fungal spores, which are produced by microorganisms. This similarity illustrates that fungal spore phenology can serve as a bioindicator of global environmental change, paralleling that of more conspicuous macroorganism phenology. Such findings underscore the potential of fungal spores to be sensitive proxies for monitoring and understanding the effects of climatic shifts on ecological systems.

### 4.2. Climate change impacts on spore season shifts

Our investigation has demonstrated significant correlations between the seasonality of fungal spores and climatic variables. We found a marked pattern of earlier spore season onset with increasing temperatures (Fig. 5), consistent with similar studies in Europe (Anees-Hill et al., 2022). There is established evidence of a strong positive correlation between spore concentrations and mean temperature (Burch & Levetin, 2002; Troutt & Levetin, 2001), suggesting a more rapid accumulation of spores in the atmosphere under global warming, possibly reaching predetermined thresholds or percentages sooner than before. This correlation aligns with mycological research indicating that key fungal life cycle events occur earlier because of global warming, such as the advancement and extension of the green season, which provides more favorable conditions for fungal growth. For example, related studies have observed earlier fungal fruiting in response to warmer temperatures (Gange et al., 2007; Kauserud et al., 2012; Klironomos et al., 1997). Other global change factors, such as plant phenology shifts and landscape change, may also contribute to this trend (Peay & Bruns, 2014). For example, the top taxa in our dataset, Cladosporiaceae, tends to gather on all kinds of plant tissues (Bensch et al., 2012), which means plant phenological shifts might cause its spore phenological shifts.

Moreover, we found that increased rainfall correlates with an earlier start and a longer duration of the allergy season, as well as higher spore concentrations (Fig. 5). This aligns with the findings of a century-long North American mycological study that suggests heightened moisture accelerates fungal fruiting (Diez et al., 2013). Although some research shows that rainfall might reduce airborne spore levels by causing a washout effect (Katial et al., 1997; Sindt et al., 2016), this perspective primarily considers the direct impact of daily precipitation on daily spore counts. In contrast, the inherent moisture from rainfall can encourage fungal growth and sporulation (Kendrick, 2017), potentially leading to elevated spore levels in the air (Ganthaler & Mayr, 2015). Thus, the net effect of increased precipitation can promote higher overall spore concentrations, despite a possible short-term negative correlation between rainfall and spore density. Our analysis extends beyond the daily impact, analyzing the relationship between total annual precipitation and annual spore metrics, which helped us highlight the impact of the precipitation regime on an interannual scale rather than the impact of rainfall on a daily scale.

While moisture is generally associated with fungal growth and sporulation, whether rainfall is associated with spore liberation may be differential by fungal species, and dependent on the morphology of the fungi. For instance, arthroconidia is a type of spore that is produced by the segmentation of fungal hyphae. *Coccidioides* form as alternative cells within hyphae, and the lysis of hyphae under dry conditions is thought to release arthroconidia (Maddy, 1957). In contrast, arthroconidia of *Blastomyces* and *Histoplasma spp.* grow as propagules of hyphae. Prior laboratory work indicated that spores of *Blastomyces* and *Histoplasma spp.* were only mobilized following wetting, suggesting a role of rainfall in detaching spores from hyphae (McDonough et al., 1976). However, higher moisture was related to a delayed ecological spore season onset and a contraction of the season. Such conditions suggest a compression of airborne spore presence around the peak period. Therefore, we propose that moist environments may lead to a concentration of spore activity within a narrower timeframe, emphasizing the complex interplay between rainfall patterns and spore dynamics.

### 4.3. Implications for ecology, public health, and climate change

The observed shifts in fungal spore phenology and seasonal intensity bear important implications for ecological systems, public health, and climate change mitigation strategies. Ecologically, fungi are crucial in processes such as decomposition, nutrient cycling, and shaping plant diseases (Dighton, 2018). Alterations in the phenology and intensity of spore distribution may lead to disruptions in these processes, with possible consequences for ecosystem function and biodiversity. An asynchrony between the phenology of fungi and the life cycles of their host plants could profoundly affect plant-fungal interactions (Song et al., 2023). Additionally, with many insects relying on fungi for nutrition (Douglas, 2009), a shift in spore availability could create cascading effects through the food web, affecting not only these insects but also the broader array of species that interact with them, including predators, parasites, and their plant hosts. Our findings, therefore, provide far-reaching implications of fungal spore season changes for ecosystems.

From a public health standpoint, the observed trends of an earlier onset of allergy seasons could affect respiratory health, particularly among vulnerable populations (Fig. 4f). Such changes can weaken current allergy management practices, as individuals may unknowingly encounter allergens outside the traditionally expected time window, leading to unforeseen, severe allergic reactions. Public health entities must therefore integrate these temporal shifts when developing awareness campaigns and advising on allergy prevention measures. Our study can serve as an alert to both patients and clinicians about the crucial role of fungal spores in allergy pathogenesis, thereby reinforcing its critical contribution to public health advocacy. Despite fungal spores being major air allergens, their study has been overshadowed by pollen research.

Evidence suggests that health impacts associated with fungal spore exposure may outweigh those linked to pollen in asthmatic individuals (Hughes et al., 2022). Notably, our research introduces a unique method by assessing allergy seasons through specific spore allergy thresholds, thereby bridging the gap between ecological conditions and public health imperatives.

Beyond the importance of fungal spores for allergies, certain species of fungi produce infectious arthroconidia that can cause severe disease in humans when inhaled. Invasive fungal infections are on the rise globally and are of particular public health concern given high mortality rates, limited therapeutics and no vaccines, and rising antifungal resistance (Fisher & Denning, 2023). Geographic expansion of many human-pathogenic fungal species has been documented, including *Coccidioides* (Mazi et al., 2023), *Cryptococcus* (Gorris et al., 2019), *Blastomyces*, and *Histoplasma spp.*, potentially associated with climate change (Nnadi & Carter, 2021). Warming temperatures may cause such pathogenic fungi to produce more spores (Frías-De-León et al., 2022). For instance, lengthened duration of spore production by *Histoplasma capsulatum* as well as *Microsporum* and *Trichophyton* genera, which cause histoplasmosis and dermatophytosis, respectively, have been observed (Frías-De-León et al., 2022; Jr, 2018; Nnadi & Carter, 2021).

While we were unable to examine spore concentration for specific pathogenic species in this study, the findings here nonetheless may help inform our understanding of the transmission of invasive fungal infections.

Ultimately, our study stands out by addressing an important gap in allergen and phenological research, specifically regarding fungal spores. While the focus has traditionally been on macroorganisms to assess the impact of climate change on biological systems, our findings underscore the sensitivity of microorganisms to climate change. This provides an important insight, as it shows that fungal spores, like larger organisms, are also responsive to climate variability. By paralleling the shifts in allergenic fungal spores with those seen in macroorganisms, we contribute to the rising evidence that supports the use of a broad spectrum of biological indicators—from the macroscopic to the microscopic—to monitor and understand the complexities of climate change. Our study underscores the need for greater attention to fungal spore research and its role in understanding climate change impacts. It also calls for increased public awareness and readiness within the health sector to address ongoing environmental shifts.

### 4.4. Limitations

While our investigation provides valuable contributions to understanding fungal spore trends amid climatic impacts, we acknowledge several limitations and identify directions for future work. First, we aggregated spore concentrations across all fungal genera to generate phenological metrics. This approach provides robust, broad-scale insights but may overlook taxon-specific responses. Different fungal taxa, such as dry- or wet-adapted fungi, are likely to respond differently to climate conditions. This taxonomic diversity can mask specific climate-driven changes in fungal communities. To address this, future studies should aim to assess trends at the genus or species level to capture more detailed responses of fungal communities to climate change. Such insights would enable more targeted ecological and public health interventions.

Second, variations in sampling protocols across National Allergy Bureau (NAB) stations introduce potential bias in our data. For example, most stations use Burkard spore traps, while others rely on Rotorod samplers, which could affect the comparability of spore concentration measurements. However, since our analysis is centered on temporal trends rather than spatial variations, these differences are unlikely to bias our findings. We also incorporated site-specific random effects into our linear mixed-effects model, which helped account for variations in sampling methods. This approach ensures that the observed trends reflect broader temporal patterns rather than localized methodological inconsistencies.

Another limitation arises from the use of linear interpolation to fill small gaps in the daily spore concentration data. This approach, while common in aerobiological studies (Anderegg et al., 2021; Gabarra et al., 2002; Picornell et al., 2021), can introduce errors, particularly during periods of high variability in spore concentrations. However, since our primary interest lies in identifying broad temporal trends in annual phenological metrics across years rather than daily fluctuations, the potential errors from linear interpolation are less likely to affect our key findings.

Additionally, the relatively sparse network of monitoring stations, combined with gaps in data at some stations, limits the geographic scope of our conclusions. Some regions of the US may be underrepresented, leading to potential gaps in understanding the regional variability of fungal spore seasonality. Expanding the network of spore monitoring stations would improve geographic representation, enhance the statistical power of future analyses, and help to capture regional differences more accurately. Moreover, some stations had shorter periods of data collection, which might reduce the ability to detect long-term trends. Sustaining long-term monitoring networks would be crucial to better capture climate-driven phenological changes in the future.

We recognize that allergy threshold determined in single clinical studies may not be generalizable due to several uncertainties detailed in the supplementary text. Nevertheless, the estimated threshold adds usefulness to translate spore concentrations to health relevance. To ensure robustness, we tested our findings of advancing spore seasons using two thresholds of 4,506 spores m^-3^ (Caillaud et al., 2018) and 6,500 spores m^-3^ (Portnoy et al., 2004), respectively (Supplementary text) and observed consistent results (Fig. S8). Future research could incorporate multiple clinical studies across various fungal taxa to refine allergy thresholds and improve the accuracy of fungal spore season characterization from a public health perspective.

Finally, the focus on annual metrics and relatively small sample size constrained the complexity of our models. Future studies that focus on intra-annual variations on fine temporal resolutions would be better equipped to explore complex impacts of climate, such as interactions and legacy effects.

## 5. Conclusions

In summary, we identify advancing fungal spore season onset, as well as decreased concentrations over the past two decades across the continental US. These trends might partly be attributed to climate change, as we found most fungal spore metrics to significantly correlate with climatic variables, including temperature and precipitation. Changes in the phenology and intensity of fungal spore season under climate change have the potential to disrupt ecological functioning and elevate allergy-related public health risks. Our findings suggest the need to further understand and adapt to responses of fungal spore season to climate change.

## CRediT authorship contribution statement

**Ruoyu Wu:** Conceptualization; formal analysis; methodology; visualization; writing – original draft; writing – review and editing. **Yiluan Song:** Conceptualization; data curation; methodology; visualization; writing – review and editing. **Jennifer R. Head:** Writing – review and editing. **Daniel S. W. Katz:** Methodology; writing – review and editing. **Kabir G. Peay:** Writing – review and editing. **Kerby Shedden:** Funding acquisition; methodology; writing – review and editing. **Kai Zhu:** Conceptualization; funding acquisition; methodology; writing – review and editing.

## Acknowledgments

RW, YS, and KZ are supported by the National Science Foundation (NSF) grants 2244711 and 2306198. RW, YS, KS, and KZ are supported by the Michigan Institute for Data and AI in Society Propelling Original Data Science grant in 2023. YS is also supported by the Eric and Wendy Schmidt AI in Science Postdoctoral Fellowship, a Schmidt Sciences program. KGP is a CIFAR Fellow in the program Fungal Kingdom: Threats and Opportunities and is supported by NSF grants 1845544, and 1926438 and Department of Energy grant DE-SC0023661. We thank Zhu Lab for constructive comments.

## Data availability

We requested fungal spore concentration data from monitoring stations associated with the National Allergy Bureau (https://pollen.aaaai.org/<u>#/</u>), with data received on Apr 25, 2023. We retrieved climatic data from Daymet (https://daymet.ornl.gov/getdata) using R package *daymetr*. Novel R Code is currently stored on Figshare (https://figshare.com/s/b492f865037e3c874993) for private peer review. Processed data and novel R code to reproduce all analyses will be permanently archived on Zenodo upon acceptance for publication.

