## Supplementary material for "Fungal spore seasons advanced across the US over two decades of climate change": Table S1, Table S2, Table S3, Table S4, Fig. S1, Fig. S2, Fig. S3, Fig. S4, Fig. S5, Fig. S6, Fig. S7, Fig. S8, Supplementary text, supplementary text

### 1. Supplementary tables

**Table S1:** Summary of available data in 55 monitoring stations associated with the National Allergy Bureau (NAB) within the continental United States that were included in the analysis of this study.

| City | State | Station | Sampling period | %raw data availability | %data availability after interpolation | %interpolated data |
| --- | --- | --- | --- | --- | --- | --- |
| Albany | NY | Certified Allergy Consultants | 2003-05-14 to 2012-10-10 | 45.3% | 50.7% | 5.5% |
| Armonk | NY | The Louis Calder Center | 2003-03-24 to 2007-10-25 | 16.3% | 44.2% | 27.9% |
| Berkeley | MO | Saint Louis County Health Department | 2003-03-03 to 2022-12-30 | 67% | 99.2% | 32.1% |
| Bismarck | ND | North Dakota Public Health Lab | 2003-04-15 to 2005-06-22 | 14.5% | 33.7% | 19.2% |
| Chelmsford | MA | Allergy and Asthma Specialists | 2004-03-30 to 2006-06-22 | 21% | 35.8% | 14.8% |
| Coeur d’Alene | ID | Asthma and Allergy of Idaho | 2016-03-17 to 2022-10-19 | 3.4% | 25.7% | 22.3% |
| College Station | TX | Scott & White Clinic | 2003-05-20 to 2011-11-17 | 5.1% | 17% | 11.9% |
| Colorado Springs | CO | Asthma & Allergy Associates, PC | 2003-03-13 to 2005-07-07 | 44.7% | 54.1% | 9.4% |
| Dayton | OH | RAPCA | 2003-03-03 to 2018-11-01 | 62.1% | 95.2% | 33.1% |
| Erie | PA | Allergy & Asthma Associates of Northwestern PA | 2003-07-15 to 2018-10-17 | 24.9% | 49% | 24% |
| Fargo | ND | Allergy & Asthma Care Center | 2003-04-13 to 2008-06-06 | 18.3% | 35.3% | 17% |
| Houston (Station 1) | TX | Allergy & ENT Associates | 2005-09-27 to 2009-10-31 | 34.9% | 71.1% | 36.2% |
| Houston (Station 2) | TX | City of Houston | 2009-02-02 to 2022-12-30 | 60.4% | 92.5% | 32% |
| Indianapolis | IN | Allergy and Asthma Specialists | 2003-03-14 to 2009-10-21 | 28.1% | 68.1% | 40% |
| Kansas City | MO | Children’s Mercy Hospital | 2003-03-12 to 2021-07-09 | 44.1% | 65.9% | 21.8% |
| Knoxville | TN | Allergy, Asthma and Immunology | 2003-03-17 to 2018-04-25 | 16% | 51.3% | 35.3% |
| La Crosse | WI | Allergy Associates of LaCrosse | 2003-03-18 to 2017-10-20 | 16.1% | 40.2% | 24.1% |
| La Jolla | CA | Erik and Ese Banck Clinical Research Center | 2005-01-03 to 2019-12-06 | 46.2% | 86.6% | 40.3% |
| Las Vegas (Station 1) | NV | Clark County Dept. of Air Quality Management | 2003-03-10 to 2010-10-28 | 33.1% | 90.4% | 57.3% |
| Las Vegas (Station 2) | NV | University of Nevada Las Vegas | 2014-08-01 to 2022-12-30 | 91.1% | 93.5% | 2.4% |
| Lexington (Station 1) | KY | University of Kentucky Asthma Allergy & Immunology | 2003-03-03 to 2018-10-01 | 33.6% | 58.3% | 24.7% |
| Lexington (Station 2) | KY | Family Allergy & Asthma - Lexington | 2012-10-30 to 2018-05-24 | 18.3% | 51.9% | 33.5% |
| Melrose Park | IL | Allergy, Sinus, and Asthma Professionals | 2004-08-13 to 2022-09-30 | 36.4% | 53.3% | 16.9% |
| Miami | FL | Florida Center For Allergy & Asthma Care | 2014-06-24 to 2020-06-15 | 32.5% | 82.7% | 50.2% |
| Milwaukee | WI | Allergy & Asthma Centers, S.C. | 2003-03-18 to 2016-10-19 | 11.8% | 53.8% | 42% |
| Monroeville | PA | Division of Allergy, Asthma and Immunology, East Suburban Pediatrics | 2004-06-15 to 2019-10-19 | 40.3% | 65.7% | 25.3% |
| Mount Laurel | NJ | Allergy and Asthma Specialty Physicians, LLC | 2003-03-17 to 2022-07-28 | 31.7% | 62.4% | 30.7% |
| New Castle | DE | Division of Air Quality, DNREC, State of Delaware | 2006-03-02 to 2017-10-30 | 23.6% | 65.3% | 41.6% |
| New Orleans | LA | Ochsner Clinic | 2007-01-01 to 2011-10-20 | 57.2% | 88.4% | 31.3% |
| Oklahoma City (Station 1) | OK | Oklahoma Allergy & Asthma Clinic, Inc. | 2003-03-04 to 2022-12-30 | 63.2% | 96.6% | 33.4% |
| Oklahoma City (Station 2) | OK | Allergy & Asthma Center | 2003-03-04 to 2017-12-29 | 47.9% | 89.5% | 41.6% |
| Olean | NY | Fred H Lewis, MD FAAAAI | 2009-05-27 to 2022-11-23 | 25.5% | 52.9% | 27.4% |
| Orange | CA | Children’s Hospital of Orange County-Pediatric Subspecialty Faculty | 2003-03-16 to 2011-07-31 | 81% | 90.9% | 9.9% |
| Oviedo | FL | Allergy & Asthma Center of East Orlando | 2005-02-02 to 2007-06-13 | 6.3% | 31.7% | 25.4% |
| Papillion | NE | The Asthma and Allergy Center, PC | 2007-04-27 to 2022-12-15 | 48.6% | 62.8% | 14.3% |
| Philadelphia | PA | The Asthma Center | 2003-03-14 to 2019-10-25 | 29.3% | 60.1% | 30.7% |
| Roseville | CA | Allergy Medical Group of the North Area | 2003-03-12 to 2018-12-15 | 12.8% | 91.8% | 79% |
| San Antonio | TX | Wilford Hall Ambulatory Surgical Center | 2011-06-18 to 2022-12-30 | 95% | 96.1% | 1.1% |
| San Jose | CA | Theodore J. Chu, M.D. | 2003-03-02 to 2022-12-18 | 12.6% | 87.8% | 75.2% |
| Santa Barbara | CA | Sansum Clinic | 2003-01-07 to 2010-11-23 | 36.4% | 84.6% | 48.2% |
| Sarasota | FL | Dr. Mary Jelks | 2003-03-03 to 2013-08-27 | 88.9% | 93.1% | 4.2% |
| Scottsdale | AZ | Mayo Clinic Arizona | 2020-08-02 to 2022-12-29 | 20% | 52% | 32% |
| Silver Spring | MD | US Army Centralized Allergen Extract Lab. | 2003-02-25 to 2022-12-29 | 55.8% | 98.8% | 43.1% |
| Springfield | MO | Springfield -Greene County Health Department | 2010-08-24 to 2018-08-20 | 40.2% | 58.9% | 18.7% |
| St. Clair Shores | MI | Lakeshore Ear Nose & Throat Center | 2003-03-03 to 2017-09-28 | 24% | 45.1% | 21.1% |
| Sylvania | OH | Allergy Clinic Ohio - Dr. Safadi | 2015-03-08 to 2022-10-15 | 50.1% | 61.1% | 10.9% |
| Tallahassee | FL | Ronald Saff, MD FAAAAI | 2003-11-13 to 2005-07-12 | 31.2% | 43.2% | 12% |
| Tampa | FL | University of South Florida | 2003-02-28 to 2022-11-02 | 23.2% | 83.7% | 60.5% |
| Tulsa (Station 1) | OK | Allergy Clinic of Tulsa | 2003-03-04 to 2022-07-06 | 38.3% | 82.5% | 44.2% |
| Tulsa (Station 2) | OK | University of Tulsa | 2003-01-01 to 2015-12-30 | 98.2% | 99.6% | 1.4% |
| Twin Falls | ID | Asthma & Allergy of Idaho | 2010-04-19 to 2022-10-05 | 16.8% | 50.1% | 33.3% |
| Vancouver | WA | AAIM Care, LLC | 2003-01-04 to 2010-11-11 | 39% | 94.9% | 55.9% |
| Waco | TX | Allergy & Asthma Care of Waco | 2003-03-10 to 2022-12-28 | 53.4% | 97.4% | 44% |
| Walnut Creek | CA | Allergy and Asthma Group of the Bay Area | 2003-03-27 to 2022-12-15 | 18.2% | 75.3% | 57.1% |
| Wilmington | DE | Nemours/A.I. Dupont Childen’s Hospital | 2011-05-02 to 2013-12-11 | 7.9% | 49% | 41.1% |

**Table S2**: Temporal trends in the relative abundance of two main taxa across all the stations. To test the temporal shifts of taxa composition, we used linear mixed-effects models of the relative abundance of the two main taxa against year. We did not observe significant changes in these two main taxa.

| **Taxa** | **Random effects** | **SD** |  |  |  |
| --- | --- | --- | --- | --- | --- |
| Unidentified | Intercept | 0.16 |  |  |  |
|  | Year | 0.000076 |  |  |  |
|  | Residual | 0.16 |  |  |  |
|  | **Fixed effects** | **Estimate** | **SE** | **t-value** | **p-value** |
|  | Intercept | 4 | 2.7 | 1.5 |  |
|  | Year | –0.0018 | 0.0013 | -1.3 | 0.18 |
| Cladosporiaceae | **Random effects** | **SD** |  |  |  |
|  | Intercept | 0.23 |  |  |  |
|  | Year | 0.000022 |  |  |  |
|  | Residual | 0.16 |  |  |  |
|  | **Fixed effects** | **Estimate** | **SE** | **t-value** | **p-value** |
|  | Intercept | 0.098 | 2.7 | 0.036 |  |
|  | Year | 0.00021 | 0.0013 | 0.15 | 0.88 |

**Table S3**: Summary of trend and observations included in the analysis for each metric. Displayed is the number of stations and observations included in the model, the slope (estimated *β*_1_ in the linear mixed-effects models) of spore metrics with log-transformation for intensity metrics, change over 2003-2022 in day for phenology metrics and % for intensity metrics (back-transformed), and the *p*-value of the mixed-effects model for each metric, including start of spore season (SOS), start of allergy season (SAS), end of spore season (EOS), end of allergy season (EAS), length of spore season (LOS), length of allergy season (LAS), amplitude concentration (Ca), peak concentration (Cp), annual integral (AIn), and allergy season integral (ASIn).

| **Metric** | **Number of stations** | **Number of observations** | **Slope** | **Change** | ***p*-value** |
| --- | --- | --- | --- | --- | --- |
| SOS | 16 | 163 | –0.5925 | –11.26d | 0.0431 |
| SAS | 31 | 372 | –1.1711 | –22.25d | 0.0088 |
| EOS | 16 | 163 | –0.0006 | –0.01d | 0.998 |
| EAS | 21 | 242 | –1.4386 | –27.33d | 0.0088 |
| LOS | 16 | 163 | 0.6108 | +11.61d | 0.1067 |
| LAS | 19 | 218 | –0.5478 | –10.41d | 0.4823 |
| ln(Ca) | 20 | 221 | –0.017 | –27.62% | 0.0118 |
| ln(Cp) | 21 | 246 | –0.0101 | –17.42% | 0.097 |
| ln(AIn) | 22 | 266 | –0.0082 | –14.46% | 0.0822 |
| ln(ASIn) | 16 | 179 | –0.0273 | –40.44% | 0.0104 |

**Table S4**: Sensitivity test on data completeness for four fungal spore season intensity metrics. Displayed is the number of observations included in the model, the slope (estimated *β*_1_ in the linear mixed-effects models) of spore metrics with log-transformation, change over 2003-2022 in % for concentrations (back-transformed), and the *p*-value of the mixed-effects model for each intensity metric, including amplitude concentration (Ca), peak concentration (Cp), annual integral (AIn), and allergy season integral (ASIn).

| **Metric** | **Data completeness** | **Number of observations** | **Slope** | **Change** | ***p*-value** |
| --- | --- | --- | --- | --- | --- |
| ln(Ca) | 100% | 161 | –0.0118 | –20.12% | 0.137 |
|  | 90% | 194 | –0.0155 | –25.54% | 0.0257 |
|  | 80% | 221 | –0.017 | –27.62% | 0.0118 |
|  | 70% | 233 | –0.0145 | –24.13% | 0.0256 |
|  | 60% | 254 | –0.0115 | –19.63% | 0.0705 |
| ln(Cp) | 100% | 162 | –0.0103 | –17.73% | 0.1548 |
|  | 90% | 213 | –0.0104 | –17.90% | 0.1007 |
|  | 80% | 246 | –0.0101 | –17.42% | 0.097 |
|  | 70% | 265 | –0.0083 | –14.54% | 0.1546 |
|  | 60% | 372 | –0.0083 | –14.57% | 0.1047 |
| ln(AIn) | 100% | 163 | –0.0067 | –11.95% | 0.2307 |
|  | 90% | 222 | –0.0075 | –13.33% | 0.1277 |
|  | 80% | 266 | –0.0082 | –14.46% | 0.0822 |
|  | 70% | 292 | –0.0122 | –20.69% | 0.0198 |
|  | 60% | 403 | –0.0069 | –12.32% | 0.1182 |
| ln(ASIn) | 100% | 122 | –0.0308 | –44.27% | 0.0284 |
|  | 90% | 151 | –0.0265 | –39.56% | 0.0245 |
|  | 80% | 179 | –0.0273 | –40.44% | 0.0104 |
|  | 70% | 183 | –0.0233 | –35.81% | 0.0286 |
|  | 60% | 198 | –0.0155 | –25.57% | 0.1288 |

### 2. Supplementary figures

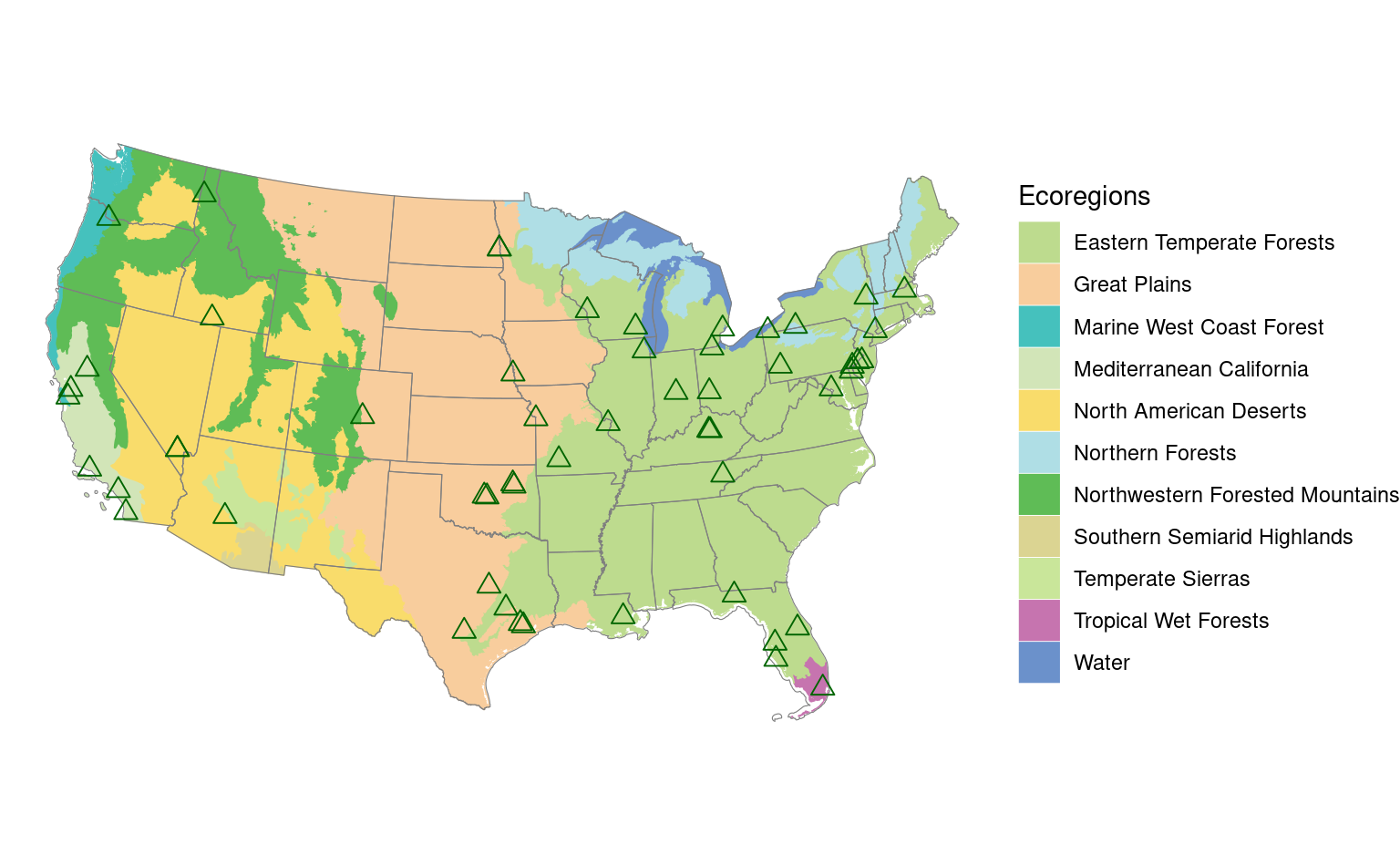

**Figure S1**: Map of 55 monitoring stations associated with the National Allergy Bureau (NAB) distributed in different ecoregions within the continental United States that were included in the analysis of this study.

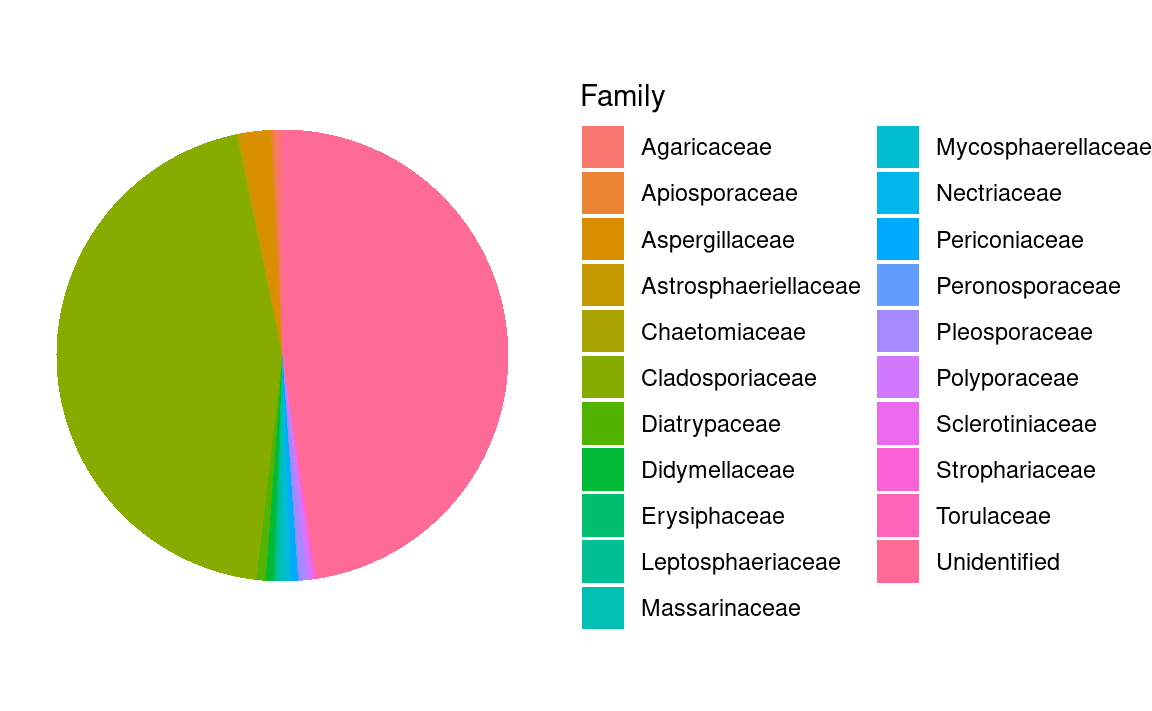

**Figure S2**: Relative abundance of 23 spore families during the study period across all the stations.

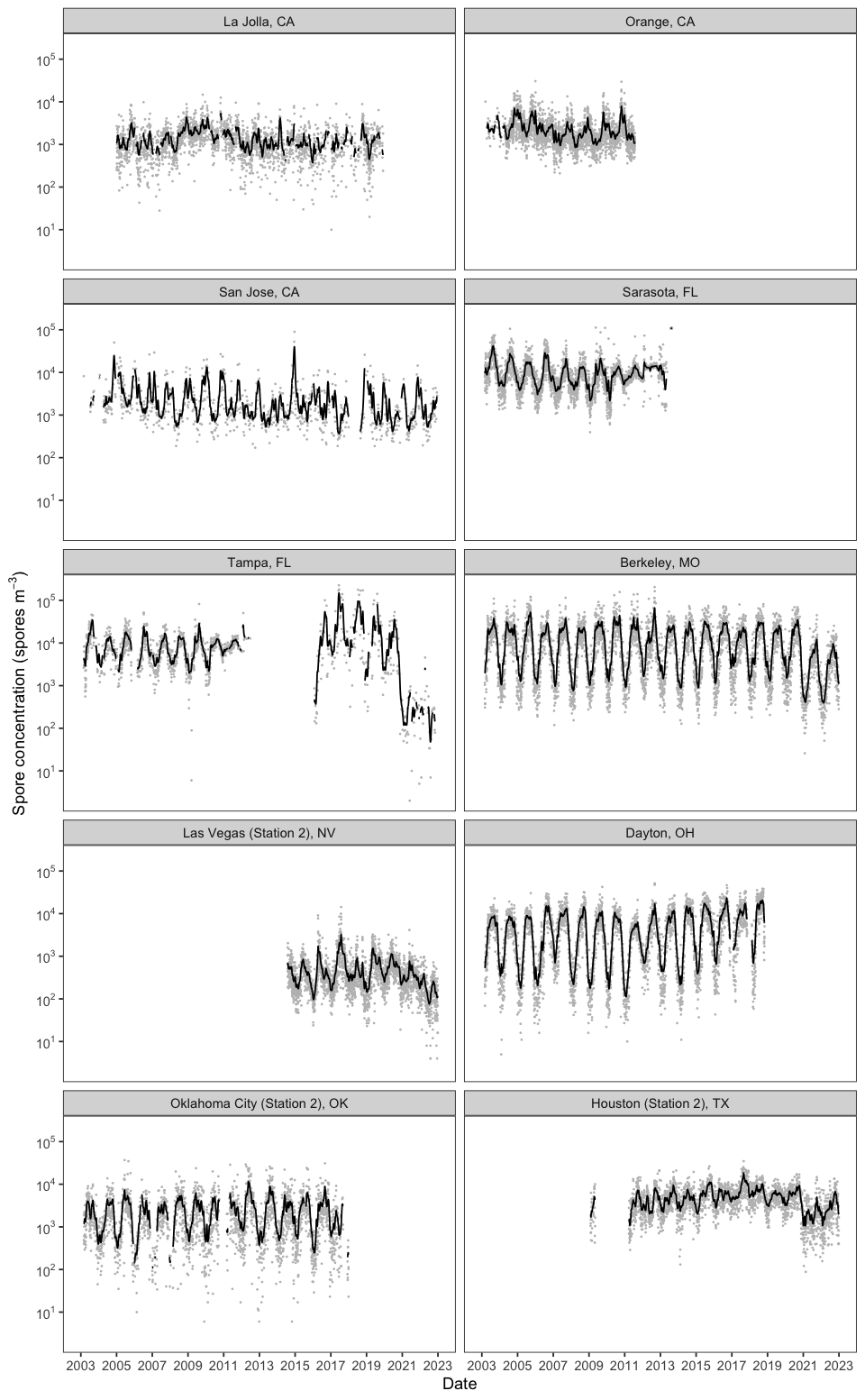

**Figure S3**: Pre-processing of raw fungal spore data. Gray dots are raw daily concentrations, while black lines are pre-processed data. Smoothing enhances the quality of the data by providing more stable and representative signals, enabling a clearer analysis of underlying trends and recurring patterns in spore season metrics. The Whittaker-Henderson smoother is particularly advantageous for its capabilities in auto-interpolating missing observations, adapting to data boundaries, and allowing for precise adjustment of smoothness levels (Eilers, 2003).

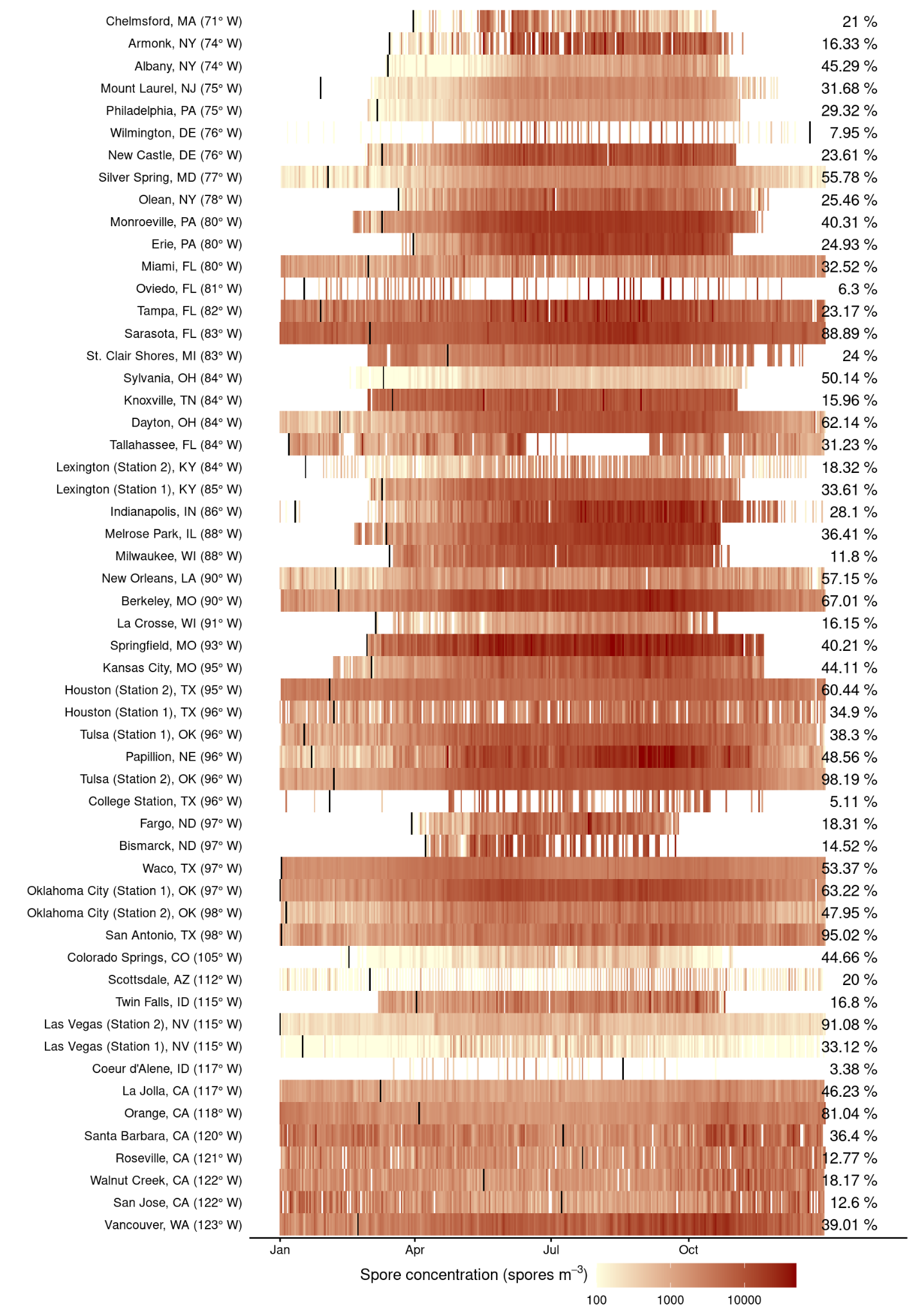

**Figure S4**: Fungal spore calendar for all the 55 stations meeting our inclusion criteria. Displayed is the daily long-term mean of fungal spore concentration, 2003-2022. Darker colors indicate higher concentrations while missing data are represented in white. Vertical black lines are the start of the spore year. Annotated numbers are the average data availability across available years for each station. Stations are ranked by longitude.

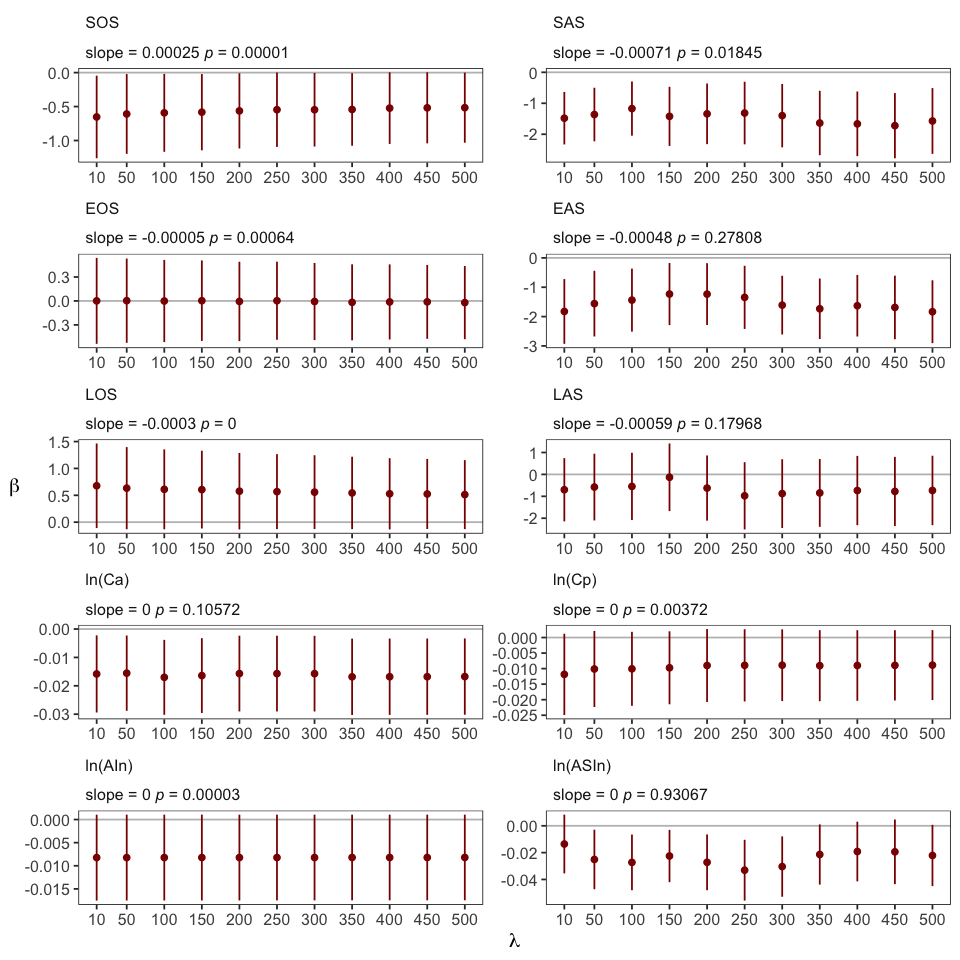

**Figure S5**: Sensitivity test on the smoothness parameter, lambda, applied in Whittaker-Henderson smoothing for ten fungal spore season metrics. The lambda was varied from ten to 500. This analysis confirmed that the observed trends in spore season were similar across the range of smoothing parameters tested. Points are estimated values of *β*_1_ in the linear mixed-effects models for trend detection. Error bars indicate the 95% confidence intervals of the fixed slope. Annotated slope and *p*-value describe the linear regression result of fixed slopes against lambda (*t*-test). Metrics defined in the ecological approach are in the left panel, while metrics defined in the public health approach are in the right panel.

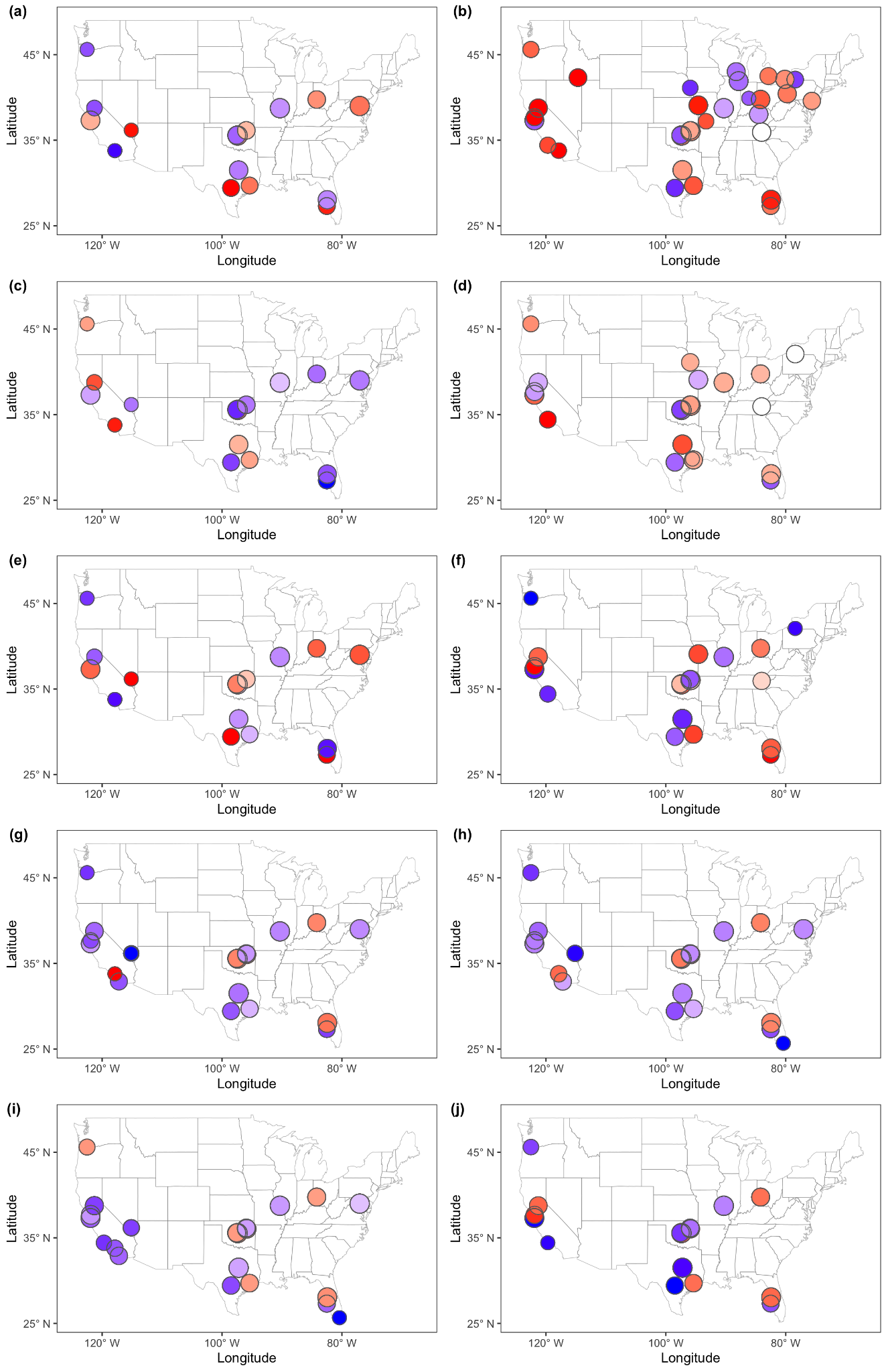

**Figure S6**: Station-level temporal trends of the spore season metrics. The trends were estimated by Theil-Sen linear regression. Redder colors indicate earlier DOY, longer days, or higher concentration, and circle sizes are proportional to the years of data at each station.

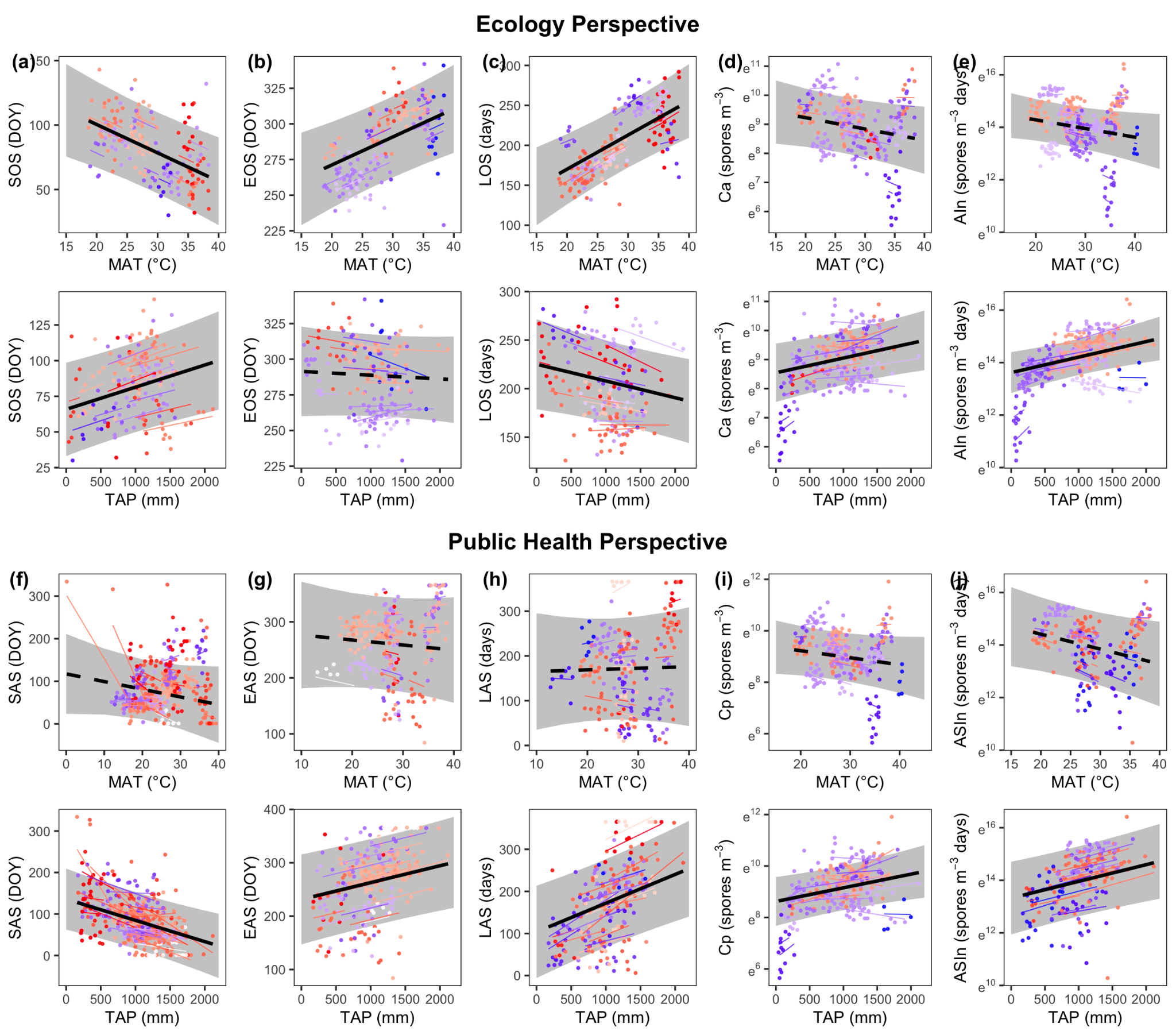

**Figure S7**: Correlation between fungal spore metrics and climate variables. Black lines are predicted slopes from linear mixed-effects models across all stations. Solid black lines indicate significant (*p* < 0.05, *t*-test) slopes. Shaded areas indicate the 95% confidence intervals of the fixed effect. Points are individual years at individual stations. Point/line colors are stations. Colors are station-level Theil-Sen linear regression slopes, with warmer colors indicating an earlier day of year, longer days, or higher intensity. The slope of colored lines are model-predicted station-level correlations. Intensity metrics are transformed using the natural logarithm.

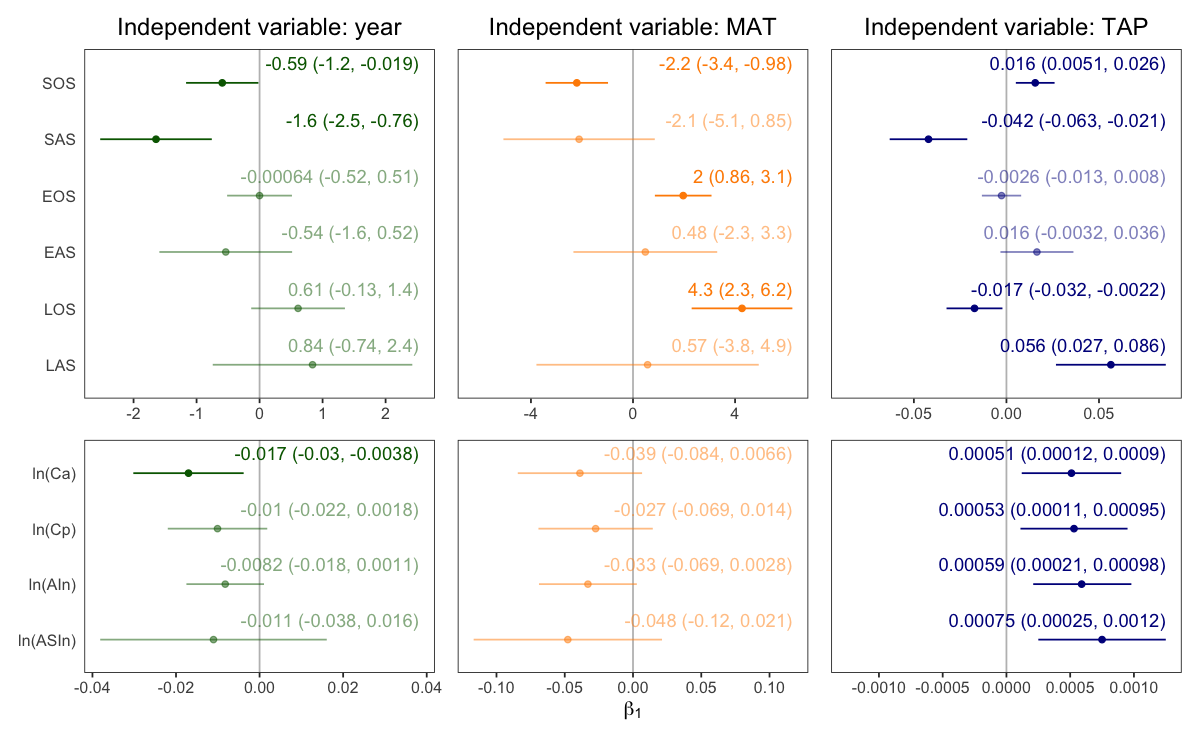

**Figure S8**: Summary of results when using NAB threshold, 6,500 spores m^–3^.

### 3. Supplementary text

In our study, we used the results of a clinical study to estimate the threshold of fungal spore concentration above which humans develop allergic reactions (Caillaud et al., 2018). From this empirical study, we obtained the relationship between outdoor mold concentration and prescribed allergy medication sales increase, expressed as the natural logarithm of the relative risk [log(RR)], which can be read as the prescribed allergy (Figure 3 of Caillaud et al., 2018). We retrieved the mold concentration at which log(RR) equals zero, representing the minimum mold concentration to induce an increase in prescribed allergy medication sales, using WebPlotDigitizer (Rohatgi, 2014). We multiplied the threshold of Cladosporium mold concentration (1,982 spores m^–3^) by the overall proportion of Cladosporium mold among all fungal spores in our dataset (44%) to estimate the threshold for total spore concentration (4,506 spores m^–3^). We then used this threshold to retrieve allergy season metrics in our analysis in the main text.

We acknowledge that there are uncertainties in this threshold for several reasons. First, we based our estimate on a study in central France. Second, we focus on the threshold of Cladosporium mold, the most dominant identified fungal spores in our dataset, without accounting for allergenic differences across fungal spore taxa. Third, there are discrepancies in dose responses between this outdoor observational study and lab experiments (Caillaud et al., 2018). Fourth, the reliance on allergy medication prescriptions as the primary outcome is limited, as it only represents a relatively indirect and attenuated response compared to more immediate health effects, such as respiratory symptoms, allergic rhinitis, or asthma exacerbations. Lastly, we did not account for differences in fungal spore composition among sites when scaling from Cladosporium mold to total spore concentration. Overall, the understudied epidemiological responses to fungal spore concentrations could lead to uncertainties in extracted allergy season metrics and long-term trends. Nevertheless, the estimated threshold serves as a starting point for understanding changing fungal spore seasons from a public health perspective.

To test the robustness of this threshold, we did the same set of analyses using a different allergy threshold reported by NAB. NAB defined four allergy thresholds: “low,” “moderate,” “high,” and “very high.” According to the NAB definitions, the moderate-level risk, set at 6,500 spores m^–3^, indicates that many individuals could be sensitive to the spores, potentially leading to allergy symptoms (Portnoy et al., 2004). We thus adopted 6,500 spores m^–3^ as the allergy threshold in our definition of allergy season to do the sensitivity test. We detected similar advanced trends in the spore and allergy season onset (Figure S8).
